# Beyond Glycolysis: Aldolase A is a Novel Effector in Reelin Mediated Dendritic Development

**DOI:** 10.1101/2024.01.12.575269

**Authors:** Gavin D. Lagani, Weiwei Lin, Sahana Natarajan, Noah Lampl, Evelyn R. Harper, Andrew Emili, Uwe Beffert, Angela Ho

## Abstract

Reelin, a secreted glycoprotein, plays a crucial role in guiding neocortical neuronal migration, dendritic outgrowth and arborization, and synaptic plasticity in the adult brain. Reelin primarily operates through the canonical lipoprotein receptors apolipoprotein E receptor 2 (Apoer2) and very low-density lipoprotein receptor (Vldlr). Reelin also engages with non-canonical receptors and unidentified co-receptors; however, the effects of which are less understood. Using high-throughput tandem mass tag LC-MS/MS-based proteomics and gene set enrichment analysis, we identified both shared and unique intracellular pathways activated by Reelin through its canonical and non-canonical signaling in primary murine neurons during dendritic growth and arborization. We observed pathway crosstalk related to regulation of cytoskeleton, neuron projection development, protein transport, and actin filament-based process. We also found enriched gene sets exclusively by the non-canonical Reelin pathway including protein translation, mRNA metabolic process and ribonucleoprotein complex biogenesis suggesting Reelin fine-tunes neuronal structure through distinct signaling pathways. A key discovery is the identification of aldolase A, a glycolytic enzyme and actin binding protein, as a novel effector of Reelin signaling. Reelin induced *de novo* translation and mobilization of aldolase A from the actin cytoskeleton. We demonstrated that aldolase A is necessary for Reelin-mediated dendrite growth and arborization in primary murine neurons and mouse brain cortical neurons. Interestingly, the function of aldolase A in dendrite development is independent of its known role in glycolysis. Altogether, our findings provide new insights into the Reelin-dependent signaling pathways and effector proteins that are crucial for actin remodeling and dendritic development.

**Significance:** Reelin is an extracellular glycoprotein and exerts its function primarily by binding to the canonical lipoprotein receptors Apoer2 and Vldlr. Reelin is best known for its role in neuronal migration during prenatal brain development. Reelin also signals through a non-canonical pathway outside of Apoer2/Vldlr; however, these receptors and signal transduction pathways are less defined. Here, we examined Reelin’s role during dendritic outgrowth in primary murine neurons and identified shared and distinct pathways activated by canonical and non-canonical Reelin signaling. We also found aldolase A as a novel effector of Reelin signaling, that functions independently of its known metabolic role, highlighting Reelin’s influence on actin dynamics and neuronal structure and growth.

## Introduction

Reelin is a secreted glycoprotein important for mammalian brain development and synaptic function (Tissir and Goffinet, 2003; Jossin, 2020). The canonical Reelin signaling pathway through the apolipoprotein E receptor 2 (Apoer2) and very low-density lipoprotein receptor (Vldlr) is best known for its role in neuronal migration and positioning in cortical layering during prenatal brain development (Caviness, 1976; Pinto-Lord et al., 1982; D’Arcangelo et al., 1995; Curran and D’Arcangelo, 1998; Trommsdorff et al., 1999). Upon Reelin binding, Apoer2/Vldlr receptor clustering triggers intracellular Dab1 phosphorylation by the Src-family tyrosine kinases (SFK) which leads to the activation of multiple downstream signaling pathways including phosphatidylinositol-3-kinase (PI3K) signaling to promote cytoskeletal changes necessary for correct neuronal positioning, axon guidance, dendrite growth and branching (Pinto Lord and Caviness, 1979; Rice et al., 1998; Hiesberger et al., 1999; Howell et al., 1999; Howell et al., 2000; Arnaud et al., 2003; Bock and Herz, 2003; Niu et al., 2004; Jossin and Goffinet, 2007; Matsuki et al., 2008; Leemhuis et al., 2010; Dillon et al., 2017). In the adult brain, Reelin signaling also promotes the maturation and stabilization of dendritic spines (Pappas et al., 2001; Niu et al., 2008; Rogers et al., 2011; Bosch et al., 2016), the primary sites of excitatory synaptic transmission and activity-dependent synaptic plasticity, where Reelin has been shown to enhance hippocampal LTP and modulate synaptic remodeling (Weeber et al., 2002; Beffert et al., 2005; Chen et al., 2005; Rogers et al., 2011; Bal et al., 2013). Therefore, a loss of Reelin or disruption in the Reelin signaling pathway has been implicated in a wide range of human neurological disorders such as lissencephaly, ataxia, autism spectrum disorders, schizophrenia, bipolar disorder, epilepsy, and Alzheimer’s disease (Impagnatiello et al., 1998; Guidotti et al., 2000; Hong et al., 2000; Perisco et al., 2001; Grayson et al., 2005).

Reelin also signals through a non-canonical pathway outside of Apoer2/Vldlr; such as integrins and ephrin receptors which act to further fine-tune Reelin-dependent migration (Dulabon et al., 2000; Bouche et al., 2013; Lee et al., 2014; Kohno et al., 2020). As such, Reelin signaling is multifaceted and activates overlapping signaling pathways following Reelin binding to its cell surface receptors. However, most of these pathways are not well characterized. Here, we investigated the role of the PI3K pathway in Reelin signaling on cytoskeleton remodeling related to dendritic outgrowth and branching. We conducted an unbiased tandem mass tag (TMT) LC-MS/MS proteomics screen to delineate canonical and non-canonical Reelin signaling and identified overlapping and distinct intracellular pathways in primary murine neurons during periods of robust neurite outgrowth. We observed pathway crosstalk between the canonical and non-canonical Reelin signaling pathways related to regulation of cytoskeleton, neuron projection development, protein transport, and actin filament-based process. We demonstrate that Reelin signaling regulates actin dynamics in developing neurites and found aldolase A, a glycolytic enzyme and actin-binding protein, as a novel downstream effector in the Reelin pathway that contributes to actin remodeling changes necessary for dendritic growth and arborization. Interestingly, the function of aldolase A on dendrite outgrowth is independent of its glycolytic function. These findings uncover novel aspects of Reelin signaling revealing broader impact beyond the well-known canonical pathways and the discovery of aldolase A as a novel Reelin effector in influencing actin remodeling and cytoskeletal dynamics on neuronal structure and growth.

## Materials and Methods

*Mice.* All animal use was approved by the Institutional Animal Care and Use Committee at Boston University and methods were performed in accordance with relevant guidelines and regulations. Timed pregnant Crl:CD1 (ICR) dams (strain code: 022, RRID:IMSR_CRL:022) were purchased from Charles River Laboratories. Mice of either sex were used for all mouse studies.

### Primary murine neuronal cultures

Primary cortical and hippocampal neurons were prepared from CD1 embryos of either sex at embryonic (E) 16.5 and E18.5, respectively. Briefly, the cortex and the hippocampi were independently dissected from each individual embryo, and dissociated with trypsin for 10 min at 37°C, triturated, and plated onto pre-treated 100 mg/mL Poly-L-lysine wells (Sigma-Aldrich Cat# P2636-100MG) 24 h prior to plating. To ensure consistency between cultures, cell density of the single cell suspension was determined using Trypan blue and a hemocytometer. Cortical neurons were plated at 7.5×10^5^ cells/well (12 well plate) and hippocampal neurons were plated at 5×10^4^ cells/well (24-well plate). Neurons were maintained in neurobasal media (Thermo Fisher Scientific Cat# 21103049) supplemented with 2% (v/v) B27 (Thermo Fisher Scientific Cat# 17504044) and 0.5 mM glutamine (Thermo Fisher Scientific Cat# 25030081) in a humidified incubator with 5% CO_2_ at 37°C.

### Generation of recombinant Reelin and GST-RAP

Stably transfected HEK293 cells (ATCC Cat# CRL-1573, RRID:CVCL_0045) expressing full-length murine Reelin were grown in T75 flasks in DMEM media and maintained at 37°C and 5% CO2. At ∼70% confluency, the media was replaced with phenol red free neurobasal media. Following 48 h, the media was collected and spun down at 500 x *g* at 4°C for 5 min to pellet cellular debris and dead cells. The supernatant was collected and concentrated using an Amicon stirred ultrafiltration cell (Millipore Cat# 5124) with a Biomax 100 kDa ultrafiltration molecular weight membrane (Millipore Cat# PBHK06210). The concentrated Reelin containing media was sterile filtered using a 0.45µm syringe filter (Millipore Cat# SLHAM33SS), aliquoted, and stored at -80°C. Mock media was generated using control vector transfected HEK293 cells and collected in parallel to the Reelin conditioned media to ensure consistency between conditions. Reelin- and mock-conditioned supernatants were tested for their ability to stimulate Reelin-dependent phosphorylation of Dab1. pGEX-RAP was transformed into BL21 DE3 *E. coli.* A single, isolated, colony was inoculated in a 5 mL starter culture and subsequently grown into 300 mL of LB media and induced with 1 mM IPTG for 4 h at 37°C. Cultures were spun down at 10,000 x *g* for 15 min at 4°C and the pellet was snap frozen in liquid nitrogen and kept at - 80°C for 15 min. Pellet was thawed and resuspended in 20 mL ice cold PBS-L (3U Benzonase/mL culture, 1% Triton X-100, 1 mM 2-Mercaptoethanol, 1 mM EDTA, 1 mg/mL Lysozyme, 2ug/mL Aprotinin, 1ug/mL Pepstatin, 1ug/mL Leupeptin in PBS). Resuspension was incubated at room temperature for 30 min on nutator and centrifuged at 4000 x *g* for 30 min at 4°C. Supernatant was run through a glutathione affinity resin column under gravity flow and GST-RAP was eluted with glutathione buffer (50 mM reduced glutathione, 50 mM Tris HCL pH 8.0 in PBS) and sterile filtered.

### Biochemical analysis and quantitative immunoblotting

For Reelin stimulation, primary murine cortical neurons were treated with 50 nM of Reelin or mock control media at 3, 7, or 14 days *in vitro* (DIV) for 30 min. GST-RAP conditions were pre-treated with 100 nM GST-RAP or GST 2 h prior to Reelin treatment. For Dab1 immunoprecipitation, neurons were lysed in immunoprecipitation buffer containing: 10 mM Tris HCL pH 8.0, 1% Triton X-100, 150 mM NaCl, 1 mM EDTA pH 8.0, 2 μg/mL Aprotinin, 1 μg/mL Pepstatin, 1 μg/mL Leupeptin, and Phosphatase inhibitor cocktail (Bimake Cat# B15001). Cell lysates were incubated with precipitating antibody against Dab1 (mouse anti-Dab1 #2721, 1:100, kindly provided by Dr. Joachim Herz) for 2 h at 4°C followed by overnight incubation with 20 μL of protein G Ultralink resin (ThermoFisher Cat# 53128). The resins were washed with immunoprecipitation buffer and precipitated proteins were eluted with reducing SDS sample buffer, boiling for 5 min and resolved by SDS-PAGE. For pathway inhibitor treatments, neurons were treated either with 50 μm LY294002 (Cell Signaling Technology Cat# 9901S), 200 nM Rapamycin (Millipore Cat# 553211), or DMSO vehicle control for 30 min prior to Reelin stimulation. Briefly, cells were rinsed with PBS and collected in RIPA buffer (150 mM NaCl, 1% NP-40, 0.5% sodium deoxycholate, 0.1% SDS, 50 mM Tris-HCL pH 8.0) supplemented with protease and phosphatase inhibitors. For Western blotting, nitrocellulose membranes were blocked in 1:1 solution of PBS and LI-COR blocking buffer for 1 h at room temperature and incubated in primary antibodies overnight at 4°C. Primary antibodies used were mouse anti-Akt (Cell Signaling Technology Cat# 2920S, 1:2000, RRID:AB_1147620), rabbit anti-Phospho-Akt Ser473 (Cell Signaling Technology Cat# 4060S, 1:2000, RRID:AB_2315049), mouse anti-p70 S6 kinase alpha (Santa Cruz Biotechnology Cat# sc-8418, 1:500, RRID:AB_628094), rabbit anti-Phospho-p70 S6 kinase Thr389 (Cell Signaling Technology Cat# 9205S, 1:1000, RRID:AB_330944), mouse anti-GAPDH (Millipore Cat# MAB374, 1:2000, RRID:AB_2107445), mouse anti-Dab1 (Santa Cruz Biotechnology Cat# sc-271136, 1:500, RRID:AB_10610240), mouse anti-Phosphotyrosine 4G10 (Millipore Cat# 05-1050, 1:1000, RRID:AB_916371). Membranes were washed and incubated in IRDye 800CW goat anti-mouse IgG (LI-COR Biosciences Cat# 926-32210, 1:20,000, RRID:AB_621842) and IRDye 680RD goat anti-rabbit IgG (LI-COR Biosciences Cat# 926-68071, 1:20,000, RRID:AB_10956166) for 1 h at room temperature and imaged using a LI-COR Biosciences Imaging Odyssey Clx System.

### Digitonin permeabilization assay

Primary murine cortical neurons were briefly rinsed with ice-cold PBS and permeabilized with 150 μL of 30 μg/mL digitonin (Sigma Cat# D141-100MG) in PBS for 5 min at 4°C. The supernatant was collected after incubation, and cells were lysed with 200 μL RIPA buffer as stated above. Equal volumes of supernatant and lysate were run on 10% Tris-glycine SDS-PAGE, transferred to nitrocellulose membrane and immunoblotted for rabbit anti-aldolase A (Proteintech Cat# 11217-1-AP, 1:2000, RRID:AB_2224626) and mouse anti-β-actin (Cell Signaling Technology Cat# 3700S, 1:2000, RRID:AB_2242334) overnight at 4°C.

### Immunocytochemistry

Primary murine hippocampal neurons cultured on 12 mm glass coverslips (Electron Microscopy Sciences Cat# 72222-01) were briefly rinsed with ice-cold PBS and fixed in 4% paraformaldehyde (PFA) for 10 min at room temperature. Cells were permeabilized in 0.1% Triton X-100 in PBS for 15 min and blocked in 5% normal goat serum in PBS for 1 h. Primary antibodies used included mouse anti-β-actin (Cell Signaling Technology Cat# 3700S, 1:1000, RRID:AB_2242334), rabbit anti-Phospho-Cofilin Ser3 (Cell Signaling Technology Cat# 3313S, 1:500, RRID:AB_2080597), or mouse anti-MAP2 (Millipore Cat# MAB3418, 1:2000, RRID:AB_11212326). To selectively stain F-actin, we use Alexa Fluor Plus 555 Phalloidin (Thermo Fisher Scientific Cat# A30106, 1:400) in 5% goat serum in PBS for 1 h at room temperature. Following PBS washes, neurons were incubated with the fluorophore-conjugated secondary antibodies goat anti-rabbit IgG Alexa Fluor-488 (Thermo Fisher Scientific Cat# A-11008, 1:500, RRID:AB_143165) and goat anti-mouse IgG Alexa Fluor-546 (Thermo Fisher Scientific Cat# A-11003, 1:500, RRID:AB_2534071) for 1 h at room temperature. Coverslips were mounted on Superfrost microscope slides (Fisher Scientific Cat# 12-550-15) in ProLong-Gold Antifade mountant with DNA stain DAPI (Fisher Scientific Cat# P36931).

### Molecular cloning and lentivirus production

The shRNA construct for mouse aldolase A was generated using oligos directed towards the 3’UTR of *Aldoa* (GCCCACTGCCAATAAACAACT) and control scrambled shRNA (CCGCAGGTATGCACGCGT) and cloned into the pLKO.1 cloning vector (Addgene Cat# 10878, RRID:Addgene_10878) and pCGLH vector for lentivirus and *in utero* electroporation experiments, respectively. To generate the lentiviral mouse R42A aldolase A mutant, site directed mutagenesis was performed with primer set (forward: CATTGCCAAGGCTCTGCAGTCCATTGG; reverse: CTTCCGGTGGACTCATCT) using the full-length mouse *Aldoa* cDNA (Sino Biological Cat# MG52539-U) and Q5 Site-Directed Mutagenesis kit (New England Biolabs Cat# E0554S). Both aldolase A wild type and R42A mutant were independently subcloned into pEGFP-C3 using primer set (forward: ACGGAATTCCAATGCCCCACCCATAC; reverse: CCAGGATCCTTAGTAGGCATGGTTAGAGATG), and subsequently subcloned into lentiviral pFUW vector using primer set (forward: ACGTCTAGAATGGTGAGCAAGGGCGAG; reverse: CCAGGATCCTTAGTAGGCATGGTTAGAGATG). To generate the lentiviral mouse D33S aldolase A mutant, site directed mutagenesis was performed with primer set (forward: GGCTGCATCTGAGTCCACCGGA; reverse: AGGATGCCCTTGCC) using the pFUW-EGFP-aldolase A wild type construct and Q5 site directed mutagenesis kit. Recombinant lentiviruses were produced by transfecting HEK293T cells (ATCC Cat# CRL 3216, RRID: CVCL 0063) with pFUW plasmids for EGFP-aldolase A wild type, EGFP-R42A aldolase A mutant, EGFP-D33S aldolase A mutant, or shRNA constructs with viral enzymes and envelope proteins (pMDLg/RRE, pRSV-REV and pVSV-G) using FuGENE6 transfection reagent (Promega Cat# E2692). HEK293T media was replaced with Neurobasal media (Thermo Fisher Scientific Cat# 21103049) supplemented with 2% (v/v) B27 (Thermo Fisher Scientific Cat# 17504044) 12 h after transfection, and the supernatant was collected 12 h after media change. Lentivirus-containing conditioned media were centrifuged at 500 x *g* for 5 min at 4°C to remove cellular debris and concentrated using Takara LentiX (Takara Cat# 631231). Briefly, the supernatant was mixed with LentiX and incubated for 30 min at 4°C followed by centrifugation at 1500 x *g* for 45 min at 4°C. The pellet was resuspended in 10% original volume in Neurobasal and stored at -80°C.

### In utero electroporation (IUE)

IUE was performed on timed pregnant Crl:CD1 (ICR) dams (strain code: 022, RRID:IMSR_CRL:022) dams at E15.5 as described previously (Gal et al., 2006; Dillon et al., 2017). Dams were anesthetized with an intraperitoneal injection of a ketamine/xylazine mixture, and the uterine horns were exposed by midline laparotomy. One to two microliters of plasmid DNA (final concentration of 1 μg/μL) mixed with 0.25% fast green dye (Sigma Aldrich Cat#F7252) was injected into the lateral ventricles using a pulled glass micropipette. For electroporation, the anode of a Tweezertrode (Harvard Apparatus) was placed over the dorsal telencephalon above the uterine muscle and four 36 mV pulses (50 ms pulse duration separated by 500 ms interval) were applied with a BTX ECM830 pulse generator (Harvard Apparatus). Following electroporation, the uterine horns were returned to the abdominal cavity and filled with warm, sterile 0.9% saline. Absorbable sutures (Havel Cat# HJ398) were used to close the abdomen and non-absorbable silk sutures (AD Surgical Cat# M-S418R19) were used to stich the skin above the abdominal wall. Dams were returned to a pre-warmed clean cage and monitored closely during recovery.

### Immunohistochemistry

Electroporated postnatal (P) day 14 mice were anesthetized with an intraperitoneal injection of ketamine/xylazine and underwent transcardial perfusion with ice cold 20 mL PBS followed by 20 mL ice cold 4% PFA. Brains were removed and fixed in 4% PFA and 5% sucrose in PBS overnight at 4°C. Brains underwent cryoprotection through a series of dehydration steps in 10%, 20%, and 30% sucrose in PBS. Brains were frozen in tissue molds with OCT compound and stored at -80°C. Coronal 40 μm brain sections were cut using a Leica CM1850 cryostat and mounted on SuperFrost microscope slides (Fisher Scientific Cat# 12-550-15) and stored at -80°C. Prior to immunostaining, sections were brought to room temperature in the dark and rehydrated using PBS and treated with 0.3% methanol peroxidase for 10 min to quench endogenous peroxidase activity. Antigen retrieval was performed by microwaving brain sections in 10 mM sodium citrate buffer pH 6.0 at 900 W for 90 s followed by 100 W for 10 min. Sections were washed in PBS and blocked in 5% normal goat serum, 0.3% Triton X-100 in PBS for 1 h at room temperature followed by incubation with rabbit anti-GFP (Synaptic Systems Cat# 132 002, 1:500, RRID:AB_887725) or mouse anti-SATB2 (Abcam Cat# ab51502, 1:200; RRID:AB_882455) and DAPI to distinguish cortical layers. in 5% goat serum PBS overnight at 4°C. Sections were washed with PBS and incubated with goat anti-rabbit IgG Alexa Fluor-488 (Thermo Fisher Scientific Cat# A-11008, 1:500, RRID:AB_143165) for 1 h at room temperature before being mounted with ProLong-Gold with DAPI.

### Image acquisition and analysis

Images were captured using a Carl Zeiss LSM700 scanning confocal microscope with image acquisition settings kept constant between coverslips in independent experiments including settings for the laser gain and offset, scanning speed, and pinhole size. To determine fluorescence intensity in neurites from primary neuron cultures, a maximum intensity projection image of the neuron was generated through imaging the entire Z-stack of the neuron at 1 µm increments with a 40X oil objective. The regions of interest were selected manually in each image using well isolated neuronal processes. Selected neurites were straightened, and the pixel intensity was quantified using National Institutes of Health ImageJ software (http://imagej.nih.gov/ij). Experimenters were blind to conditions during image acquisition and analysis. For Sholl analysis, healthy, non-overlapping neurons were chosen to minimize neurite crossover. Sholl analysis was conducted using the Neuroanatomy and NeuronJ plugins for ImageJ (Meijering et al., 2004). Concentric circles were spaced 10 µm apart starting from the center of the soma and extending to the end of the longest neurite. For Sholl analysis from IUE experiments, isolated GFP cell-filled neurons in primary somatosensory cortex (S1) were chosen. Z-stack images of selected neurons were acquired using a 25X oil objective and a 0.5 µm interval step. Full Z-stack images were 3D reconstructed using IMARIS image analysis software, and neuronal processes were manually traced to highlight the neuronal arborization pattern. Sholl analysis was conducted using concentric circles spaced 1μm apart starting from the center of the soma and going to the end of the longest traced neurite. For apical neurite orientation, well isolated neurons in layers 2/3 of S1 were chosen. Maximum intensity projection images were generated from Z-stack images of neurons taken with a 1 µm step interval using a 25X oil objective. Apical neurite orientation was determined using NIH ImageJ. A perpendicular line was drawn from the center of the soma to the pia and a second line following the trajectory of the apical neurite was drawn starting at the soma. The angle between the two lines was used to determine neurite angle to the pia. Apical neurites were identified as the longest neurites originating at the apex of the soma with a trajectory of ≤ 90° to the pia. For layering and migration analysis, single plane images of cortical columns spanning the pia to the white matter in S1 were captured using a 25X oil objective. Using NIH ImageJ, boundaries between cortical layers were denoted, and the number of GFP positive somas were counted in each layer. To determine average distance from the pia, a perpendicular line to the pia from the center of GFP labeled somas was drawn using NIH ImageJ.

### Liquid chromatography tandem mass spectrometry

Primary murine neurons were plated at 2×10^6^ cells/well in 6 well plates and treated with 50 nM Reelin, mock media control for 30 min or 100 nM GST-RAP for 2 hours prior to Reelin treatment at DIV 7. The media was aspirated and cells were briefly rinsed in ice-cold PBS and collected in 750 μL of ice-cold PBS and centrifuged at 500 x *g* for 2 min at 4°C. Cell pellets were snap frozen in liquid nitrogen and stored at -80°C. Samples were resuspended in 200 μL of GuHCL lysis buffer (6 M GuHCL, 100 mM Tris pH 8.0, 40 mM chloroacetamide, 10 mM TCEP) with phosphatase inhibitor cocktail (Roche Cat# 4906837001) and EDTA free protease inhibitor cocktail (Thermo Fisher Scientific Cat# P190058) followed by heating to 95°C for 10 min. After brief sonication using a Branson probe sonicator with 10% power (40 KHz) for 10 s on ice to shear DNA, the samples were quantified *via* BCA assay (Thermo Fisher Scientific Cat# 23225). The samples were diluted with 100 mM Tris, pH 8.5 buffer to bring GuHCL concentration down to 0.75M. Lysate proteins were digested with LC/MS-grade trypsin (1:50 enzyme to protein ratio, w/w) at 37°C overnight with shaking followed by the addition of formic acid (FA) to 1% in solution to terminate the trypsinization. The resulting peptides were desalted using a C18 solid phase extraction Sep-Pak column (SPE, Waters) as per the manufacturer’s instructions. Briefly, after SPE columns were activated using 90% methanol and pre-conditioned using 0.1% TFA, the peptide digests were loaded, washed with buffer of 0.1% TFA for 2 times and eluted with 0.1% TFA-60% acetonitrile. The desalted peptides were dried under vacuum at 45 °C and kept at -20°C prior to tandem mass tags (TMT) labeling. Prior TMT labeling, peptide quantification was performed by Pierce quantitative colorimetric assay (Thermo Fisher Cat# P123275). Each sample comprising 100 μbated with TMTPro 16plex reagents (Thermo Fisher Scientific Cat# A44520) as per the manufacturer’s protocol. Reaction was carried out for 1 h at room temperature. To quench the reaction, 5% hydroxylamine was added to each sample and incubated for 15 min. Equal amounts of each sample were combined in a new tube and a speed vac was used to dry the labelled peptide sample. The labelled peptides were desalted using a C18 SPE column. TMT-labeled peptides were then fractionated offline on a Waters XBridge BEH C18 reversed-mL/min with two buffer lines: buffer A (consisting of 0.1% ammonium hydroxide-2% acetonitrile-water) and buffer B (consisting of 0.1% ammonium hydroxide-98% acetonitrile, pH 9). The peptides were separated by a gradient from 0% to 10% B in 5 min followed by linear increases to 30% B in 23 min, to 60% B in 7 min, and then 100% in 8 min and maintained at 100% for 5 min. This separation yielded 48 collected fractions that were subsequently combined into 12 fractions and evaporated to dryness in a vacuum concentrator. Fractions were kept at -20°C prior to LC-MS analysis. LC-MS analysis was performed using an Orbitrap Exploris 480 mass spectrometer equipped with FAIMS and interfaced to an Easy nanoLC1200 system (Thermo Fisher Scientific Cat# LC140). Peptides were loaded onto a C18 pre-column (3 μm, 75 mm i.d. × 2 cm, 100 Å, Thermo Fisher Scientific Cat# 16-494-6) then separated on a reverse-phase nano-spray column (2 μm, 75 mm i.d. × 50 cm, 100 Å, Thermo Fisher Scientific Cat# 16-494-6) using gradient elution. Samples (approximately 2 μg peptides) were injected and separated over 150 min gradient. The mobile phase A was consisted of 0.1% FA-2% acetonitrile, and mobile phase B was consisted of 0.1% FA-80% acetonitrile. The gradient consisted of 5% to 30% mobile phase B over 100 min, was increased to 60% mobile phase B over 20 min then was increased to 99% mobile phase B over 2 min and maintained at 95% mobile phase B for 5 min at a flow rate of 250 nL/min. The mass spectrometry of 60,000 with a normalized AGC target of 300% for peptide. The source ion transfer tube temperature was set at 275°C and a spray voltage set to 2.5kv. Data was acquired on a data dependent mode with FAIMS running 3 compensation voltages at -50v, -57v and -64v. MS2 scans were performed at 45,000 resolution with a maximum injection time of 80 ms for peptides at normalized collision energy 34. Dynamic exclusion was enabled using a time window of 60 s.

### Proteomic data analysis

MS2 spectra were processed and searched by MaxQuant (version 1.6) against a database containing the Uniprot mouse protein sequences (uniprot.org) and reversed (decoy) sequences for protein identification. The search allowed for two missed trypsin cleavage sites, variable modifications of methionine oxidation, and N-terminal acetylation. The carbamidomethylation of cysteine residues was set as a fixed modification. Ion tolerances of 20 and 6 ppm were set for the first and second searches, respectively. The candidate peptide identifications were filtered assuming a 1% false discovery rate (FDR) threshold based on searching the reverse sequence database. Quantification was performed using the TMT reporter on MS2 (TMT pro16-plex). Bioinformatic analysis was performed in the R (version 4.3) statistical computing environment. Relative intensity of protein groups features was log2 transformed then normalized by median method. The Limma R package was used for differential analysis (Ritchie et al., 2015) to generate ranked lists. The differential analysis and comparison between groups was performed using a Student *t*-test. Fast Gene Set Enrichment Analysis (fgsea) (Korotkevich et al., 2021) with statistical significance calculated using 10,000 permutations was used to identify biological terms, pathways and processes that were co-ordinately up- and down-regulated within each pairwise comparison. FDR cut-off was set to 0.05 for significant terms. Bar-plots were generated with the ggplot Rpackage using the normalized enrichment score (NES). The exact p-value, FDR, and NES of GSEA enrichment were included in supplementary table. The mass spectrometry proteomics raw data have been deposited to the ProteomeXchange Consortium via the PRIDE repository with the dataset identifier PXD046667.

### Quantification and data analysis

Experimenters were blind to condition during data acquisition and analysis. Statistical parameters are presented as mean ± S.E.M. To determine statistical significance between groups, unpaired *t*-test was used to analyze all pairwise datasets and one or two-way analysis of variance (ANOVA) for parametric analysis of multiple comparisons followed by Tukey’s or Šídák’s *post hoc* test. To determine statistical significance of the Sholl analysis, a two-way ANOVA was used. Analysis was conducted using Prism 8 software (GraphPad Software, RRID:SCR_000306). Significance was set to *p* < 0.05 for all experiments. Number of experiments and statistical information are reported on the corresponding result and figure legend sections.

## Results

### Reelin differentially activates PI3K/Akt and S6K1 phosphorylation across neuron development

We and others have shown when purified recombinant Reelin is added to wild type murine cultured neurons, Reelin stimulates the phosphorylation of Dab1 and PI3K/Akt signaling pathways at DIV 3 (Beffert et al., 2002; Jossin and Goffinet, 2007; Leemhuis et al., 2010). Because Reelin signaling activates overlapping signaling pathways that govern important neuronal development and functions, including synaptic transmission, it is unclear whether Reelin activates the PI3K/Akt pathways uniformly throughout neuronal and synaptic development. To address this, we first stimulated primary wild type murine neurons with 50 nM Reelin or mock control at three developmental time points, DIV 3 (axon outgrowth), DIV 7 (dendritic outgrowth) and DIV 14 (mature synapses) for 30 min and performed Dab1 immunoprecipitation to monitor Dab1 phosphorylation activity by immunoblotting with an anti-phosphotyrosine antibody. We demonstrated Reelin consistently increased the levels of phosphorylated Dab1 in immunoprecipitates (lane 2, Fig. 1*A-C*) compared to mock medium (lane 1, Fig. 1*A-C*) at DIV 3, 7 and 14.

**Figure 1.**
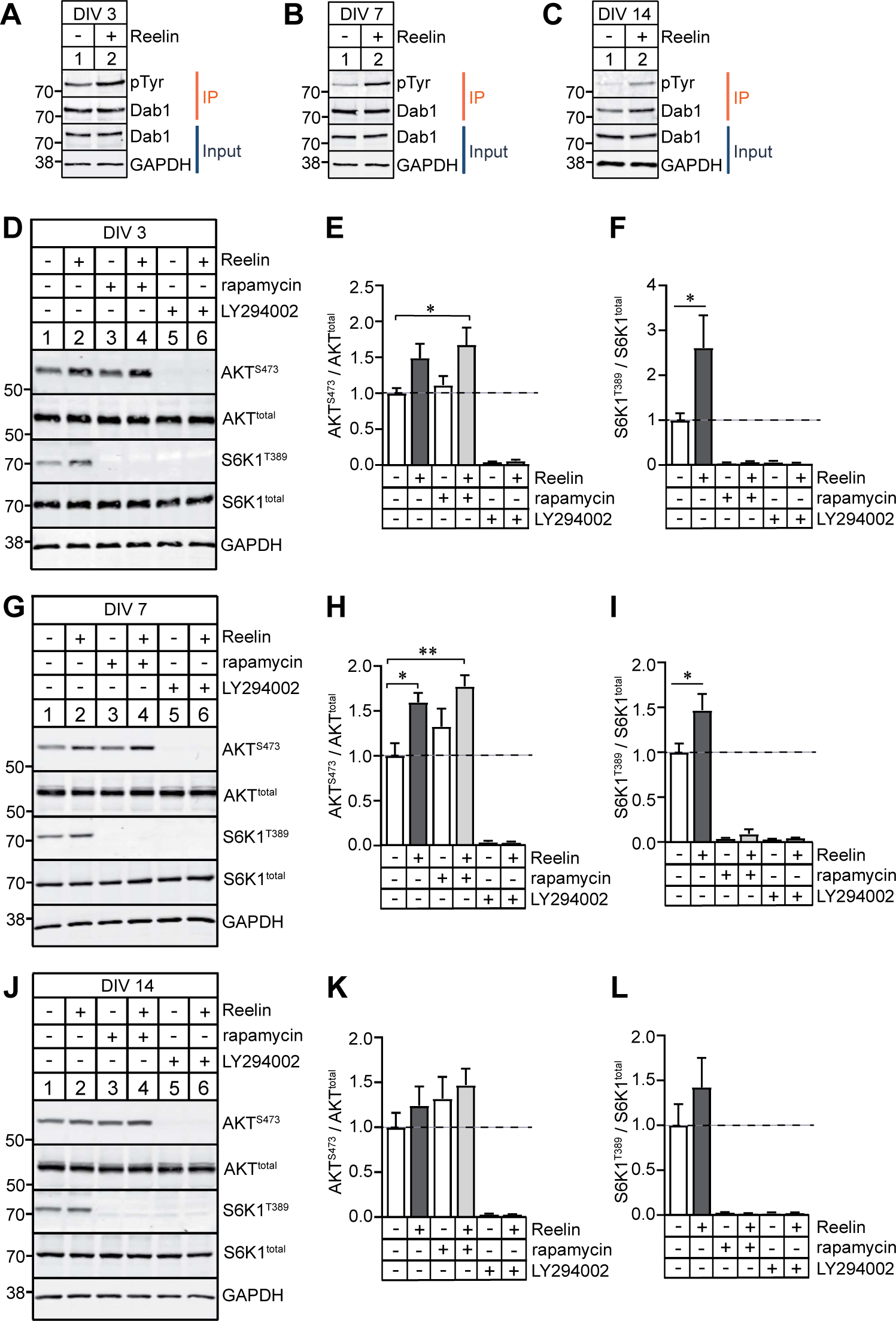
Reelin selectively activates the PI3K/Akt and S6K1 pathway in primary murine neurons at early developmental time points. ***A-C,*** Representative immunoblots showing phospho-tyrosine levels after Dab1 immunoprecipitation (IP) from primary murine neurons treated with 50 nM Reelin or mock control for 30 min at DIV 3, 7, and 14. Total Dab1 and GAPDH serve as loading controls (input). ***D,*** Representative immunoblots from DIV 3 neuronal lysates stimulated for 30 min with mock control (lanes 1, 3, 5) or with 50 nM Reelin (lanes 2, 4, 6). DIV 3 neuronal cultures were also incubated with 200 nM rapamycin (lanes 3, 4) or 50 μM LY294002 (lanes 5, 6) 30 min prior to Reelin stimulation. Cell lysates were immunoblotted for phosphorylated S473 Akt, total Akt, phosphorylated T389 S6K1, total S6K1, and GAPDH that serves as loading control. ***E,*** Bar graph quantification for the ratio of phosphorylated S473 Akt with total Akt from ***D*** shows an increase in Akt phosphorylation following Reelin stimulation that was inhibited by LY294002 [one-way ANOVA, F (5, 12) = 24.55, *p* < 0.0001]. ***F,*** Bar graph quantification for the ratio of phosphorylated T389 S6K1 with total S6K1 from ***D*** shows an increase in S6K1 phosphorylation following Reelin stimulation that was inhibited by rapamycin and LY294002 [one-way ANOVA, F (5, 12) = 11.71, *p* = 0.0003]. ***G,*** Representative immunoblots from DIV7 neuronal lysates with the same treatment conditions as described in ***D***. ***H,*** Bar graph quantification for the ratio of phosphorylated S473 Akt with total Akt from ***G*** [one-way ANOVA, F (5, 12) = 39.74, *p* < 0.0001]. ***I,*** Bar graph quantification for the ratio of phosphorylated T389 S6K1 with total S6K1 from ***G*** [one-way ANOVA, F (5, 12) = 52.55, *p* < 0.0001]. ***J,*** Representative immunoblots from DIV 14 neuronal lysates with the same treatment conditions as described in ***D***. ***K,*** Bar graph quantification for the ratio of phosphorylated S473 Akt with total Akt from ***J*** [one-way ANOVA, F (5, 12) = 15.41, *p* < 0.0001]. ***L,*** Bar graph quantification for the ratio of phosphorylated T389 S6K1 with total S6K1 from ***J*** [one-way ANOVA, F (5, 12) = 14.42, *p* = 0.0001]. Quantifications are from n = 3 individual replicates for each treatment condition and time point.

Next, we examined whether Reelin induces phosphorylation of Akt (Ser473) and S6K1 (Thr389) across all three neurodevelopmental time points. At DIV 3, we found a noticeable, although not significant, 49% increase in Akt phosphorylation following Reelin stimulation (lane 2, Fig. 1*D-E*) compared to mock control (lane 1, Fig. 1*D-E*). The 67% increase in Akt phosphorylation compared to mock control was insensitive to the downstream mTORC1 inhibitor rapamycin when neurons were treated 30 min prior to Reelin stimulation (*p* = 0.0485, lane 4, Fig. 1*D-E*). As expected, Akt phosphorylation was prevented by inhibition of upstream PI3K with LY294002 (lanes 5 and 6, Fig. 1*D-E*). In addition, we observed a 160% increase in S6K1 phosphorylation after Reelin stimulation when compared to mock control (*p* = 0.0254, lane 2, Fig. 2*D, F*) at DIV 3, and this effect was inhibited by both rapamycin and LY294002 (lanes 3-6, Fig. 1*D, F*). Likewise, at DIV 7, we observed Akt phosphorylation was increased by 59% after Reelin stimulation compared to mock control (*p* = 0.0412, lane 2, Fig. 1*G, H*) and increased to 77% in the presence of rapamycin (*p* = 0.0071, lane 4, Fig. 1*G, H*), but was inhibited by PI3K inhibitor LY294002. Reelin stimulation led to a 46% increase in S6K1 phosphorylation when compared to mock control at DIV 7 (*p* = 0.0232, lane 2, Fig. 1*G, I*). By DIV 14, we did not observe any changes in both Akt and S6K1 phosphorylation following Reelin stimulation compared to mock control (Fig. 1*J-L*). These results indicate that Reelin selectively activates S6K1 phosphorylation through the PI3K/Akt and the mTORC1 complex pathway during early axonal and dendritic outgrowth but not in mature neurons.

**Figure 2.**
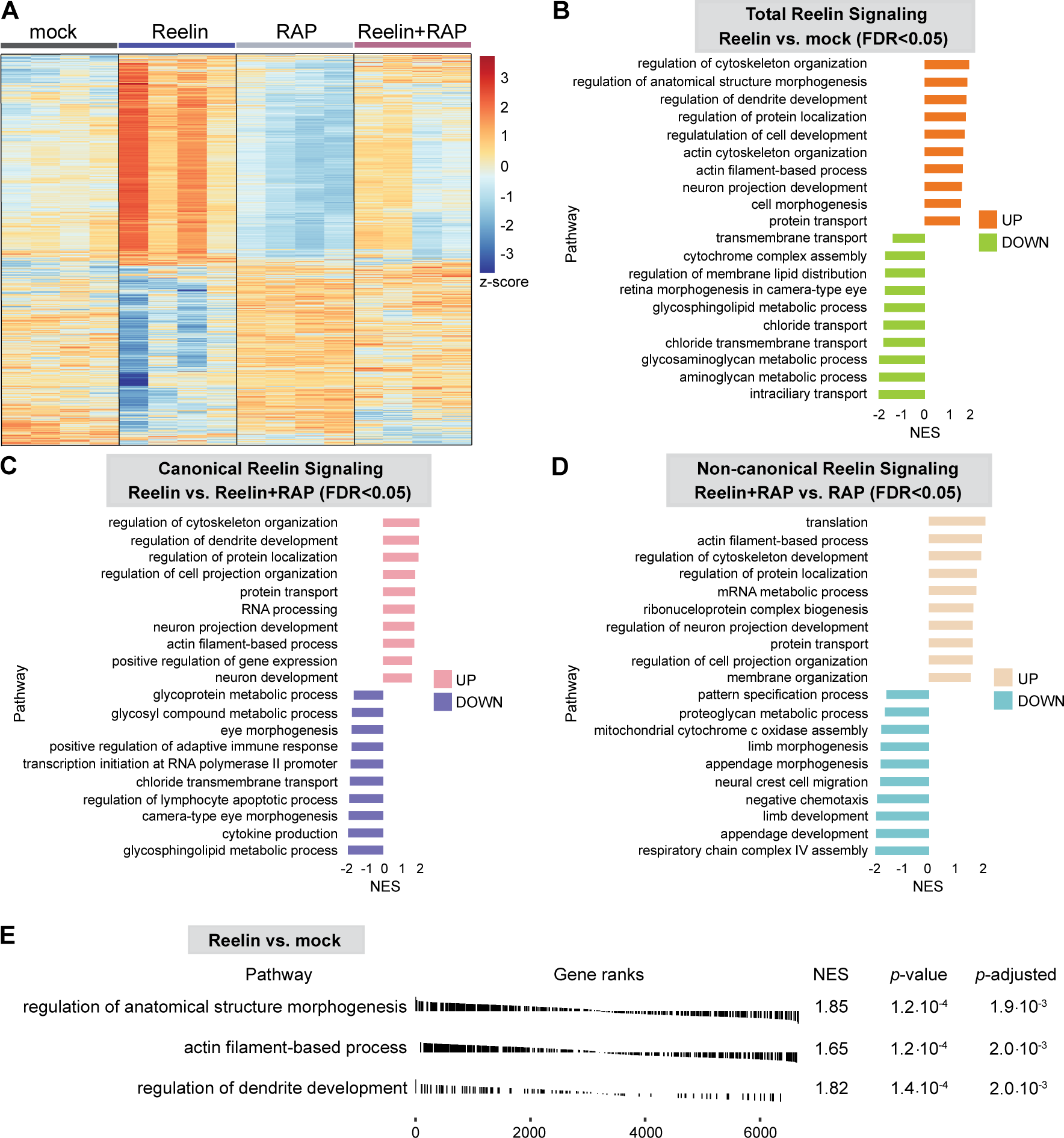
Proteomic analysis of acute Reelin signaling. Primary murine neurons at DIV 7 were treated with mock control, 50 nM Reelin for 30 min, 100 nM GST-RAP for 2 h, or Reelin with GST-RAP (2 h prior to 30 min Reelin stimulation) and subject to TMT LC-MS/MS. ***A,*** Heatmap of proteome clusters indicating differentially-expressed proteins (rows) that are significantly upregulated (red) and downregulated (blue). ***B,*** Two-side bar plot of GSEA pathways identified from the Reelin and mock control comparison with FDR < 0.05. The numbers at the bottom are normalized enrichment scores (NES) for the corresponding biological pathway categories that are up- and down-regulated of the ranked list. ***C,*** Two-side bar plot of GSEA pathways identified from the Reelin and Reelin with RAP comparison with FDR < 0.05 representing the canonical Reelin pathway. ***D,*** Two-side bar plot of GSEA pathways identified from the Reelin with RAP and RAP alone comparison with FDR < 0.05 representing the non-canonical Reelin pathway. ***E,*** Gene ranking plot of several developmental and cytoskeletal associated pathways enriched in Reelin stimulation compared to mock control, *p* ≤ 0.02.

### Gene set enrichment analysis support roles of Reelin in regulation of cytoskeleton development and organization particularly related to actin filament-based process

Since we established Reelin activates the PI3K/Akt and mTORC1 signaling pathway during early neuronal development, we next wanted to elucidate the proteomic changes induced by Reelin stimulation during dendritic outgrowth at DIV 7, a period where we observed consistent PI3K/Akt and S6K1 phosphorylation. A preliminary time course in primary murine neurons at DIV 7 demonstrated Reelin exerted a prominent effect on the proteome at 30 min compared to 3, 6 or 24 h using Label Free Quantification Mass Spectrometry (LFQ-MS) (Supplementary Fig. 1). To examine the proteomic changes more extensively at 30 min, we treated DIV 7 primary murine neurons in quadruplicates with four different conditions: mock control, 50 nM Reelin, 100 nM GST-RAP, or Reelin and GST-RAP and performed liquid chromatography tandem mass spectrometry (LC-MS/MS) analysis. RAP is a chaperone protein that acts in the secretory pathway to prevent premature binding of ligands to lipoprotein receptors prior to their insertion into the cell membrane (Bu and Rennke, 1996; Willnow et al., 1996). As such, RAP inhibits the canonical Reelin signaling pathway by preventing Reelin binding to Apoer2 and Vldlr and blocking the Reelin-induced phosphorylation of Dab1 in primary neurons (Hiesberger et al., 1999). Indeed, we demonstrated that GST-RAP effectively blocked Reelin-induced Dab1 phosphorylation when primary murine neurons were treated with 100 nM GST-RAP 2 h prior to Reelin stimulation at DIV 7 (lane 3, Supplementary Fig. 2B). With this experimental design, we can detect the following proteomic changes induced by Reelin: (a) Canonical Reelin signaling by comparing the Reelin-treated group with the Reelin and RAP-treated group where the difference represents the effect of Reelin signaling *via* the canonical receptors, Apoer2 and Vldlr which are inhibited by RAP. (b) Non-canonical Reelin signaling by comparing the Reelin and RAP-treated group with RAP alone where both groups are inhibited from interacting with the canonical receptors by RAP, so any difference between these groups would primarily be due to signaling *via* non-canonical receptors. (c) Total Reelin signaling that includes both canonical and non-canonical signaling by comparing the Reelin-treated group with mock control group where the difference represents the total effect of Reelin regardless of whether the signaling is by canonical or non-canonical receptors.

Using a high-throughput multiplex tandem mass tag (TMT) labeling approach, we identified 6744 total proteins with a stringent false discovery rate (FDR) < 0.01 across the four conditions. Reelin stimulation led to significant changes within the proteome when compared to mock control after 30 min (Fig. 2*A*). In contrast, GST-RAP treatment regulated protein expression in the opposite direction compared to the Reelin group (*p* < 0.05, Fig. 2*A*), likely due to RAP’s effects on blocking Reelin binding to Apoer2 and Vldlr. Interestingly, we observed a distinct expression pattern of proteomic changes in the Reelin and GST-RAP treatment condition indicating Reelin signaling through the non-canonical pathway has a substantial and distinct impact on the proteome.

To examine how the canonical and non-canonical Reelin pathways differ functionally, we performed gene set enrichment analysis (GSEA) on the TMT dataset. We ranked all genes according to the extent of their differential expression between the groups and computed normalized enrichment scores (NES) for the collection of gene sets representing biological pathways and identified gene sets that are up- and down-regulated of the ranked list. When we compared the Reelin-treated group with mock control where the difference represents the total effect of Reelin, we found upregulated pathways related to regulation of cytoskeleton organization, anatomical structure morphogenesis, dendrite development, protein localization and transport, cell development and morphogenesis, actin cytoskeleton organization, actin filament-based process, and neuron projection development (FDR < 0.05, Fig. 2*B*). In contrast, down-regulated genes belonged to gene sets associated with intraciliary transport, aminoglycan, glycosaminoglycan and glycosphingolipid metabolic process, and chloride transport (Fig. 2*B*). Interestingly, a majority of the upregulated pathways in total Reelin signaling overlapped with canonical Reelin signaling when we compared the Reelin-treated group with the Reelin and RAP-treated group including regulation of cytoskeleton organization, dendrite development, protein localization and transport, neuron projection development and actin filament-based process (Fig. 2*C*). This also coincided with downregulated pathways including glycosphingolipid metabolic process and chloride transmembrane transport. This was not surprising given the preferential binding of Reelin to the lipoprotein receptors Apoer2 and Vldlr. GSEA also identified gene sets unique to the canonical Reelin signaling pathway, including the upregulation of RNA processing and positive regulation of gene expression and the downregulation of cytokine production and regulation of lymphocyte apoptotic process. When we examined non-canonical Reelin signaling by comparing the Reelin and RAP-treated group with RAP alone, we identified several significantly enriched gene sets unique to the non-canonical Reelin pathway, including translation, mRNA metabolic process, ribonucleoprotein complex biogenesis. We noted several downregulated pathways related to respiratory chain complex IV assembly, appendage and limb development and morphogenesis. We also observed pathway crosstalk between the canonical and non-canonical Reelin signaling pathways related to regulation of cytoskeleton and neuron projection development, protein transport and actin filament-based process (Fig. 2*D-E*). Together, these data show that Reelin stimulation regulates proteomic networks associated with regulation of anatomical structure morphogenesis and dendrite development, particularly related to actin filament-based process at DIV 7 (Fig. 2*E*).

### Reelin stimulation regulates actin dynamics in developing neurites

Because we observed an enrichment of gene sets related to regulation of cytoskeleton development and organization, including actin filament-based process following Reelin stimulation, we explored whether Reelin functionally plays a role in actin filament dynamics in developing neurons. To test this, we stimulated DIV 7 cultured neurons with 50 nM Reelin and performed labeling for phalloidin to probe for filamentous actin (F-actin), a key cytoskeletal component important for cell motility and growth. Reelin stimulation led to a 23% decrease of phalloidin intensity in neurites when compared to mock control (*p* < 0.0001, Fig. 3*A, B*) indicating that Reelin-treated neurons may lead to F-actin remodeling in neurites. To test whether Reelin’s effects are dependent on the canonical Reelin signaling pathway, we treated neurons with 100 nM GST-RAP for 2 h prior to Reelin stimulation and found that RAP blocks the Reelin mediated decrease in phalloidin intensity. Similarly, treating neurons with PI3K inhibitor LY294002 prevented Reelin mediated effects on phalloidin intensity indicating the Reelin mediated changes in phalloidin intensity are through the canonical Reelin signaling that requires PI3K activation. We next systematically examined whether the decrease in phalloidin intensity following Reelin stimulation was localized to primary or secondary neurites. We observed Reelin stimulation led to a 24% decrease in phalloidin intensity in primary neurites when compared to mock control (*p* = 0.0012, Fig. 3*C*). In addition, we observed phalloidin intensity in the Reelin stimulated group was 26.7% lower when we compared to the Reelin and LY294002-treated group in secondary neurites (*p* = 0.0321 Fig. 3*D*).

**Figure 3.**
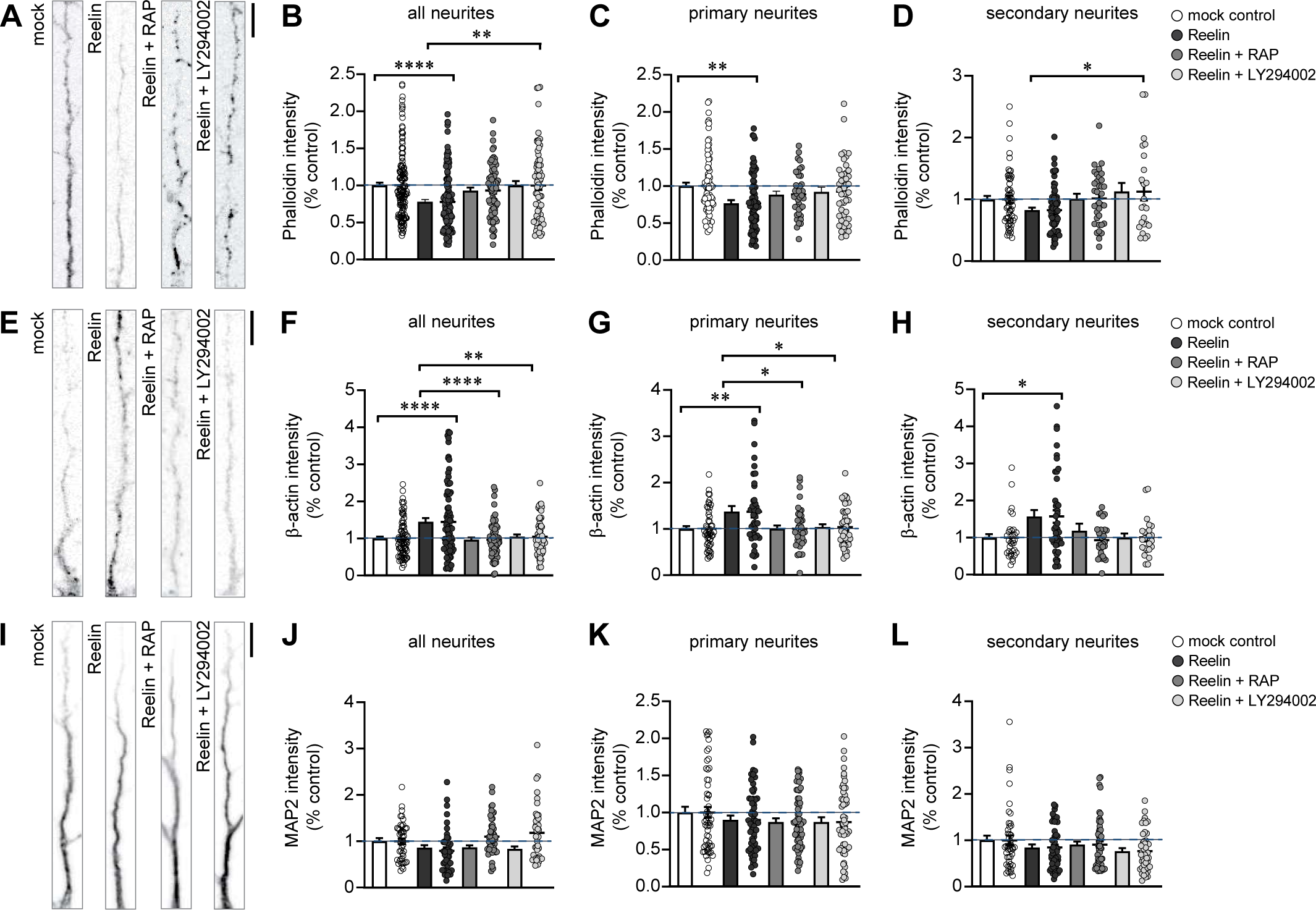
Reelin stimulation regulates actin remodeling in developing neurites. Primary murine neurons at DIV 7 were treated with mock control, 50 nM Reelin, 100 nM GST-RAP for 2 h prior to Reelin or 50 μM LY294002 for 30 min prior to Reelin stimulation (30 min). ***A,*** Representative images of neurons labeled with phalloidin showed a decrease in phalloidin intensity in neurites following Reelin stimulation. ***B,*** Bar graph quantification of fluorescent phalloidin intensity in all neurites normalized to mock control [one-way ANOVA, F (3, 423) = 7.879, *p* < 0.0001]. ***C,*** Bar graph quantification of phalloidin intensity in primary neurites normalized to mock control [one-way ANOVA, F (3, 239) = 4.820, *p* = 0.0028]. ***D,*** Bar graph quantification of phalloidin intensity in secondary neurites normalized to mock control [one-way ANOVA, F (3, 180) = 3.307, p = 0.0214]. n = 5 independent experiments with n = 15-26 neurons per condition analyzed. ***E,*** Representative images immunostained for beta-actin (β-actin) showed an increase in β-actin intensity in neurites following Reelin stimulation. ***F,*** Bar graph quantification of fluorescent β-actin intensity in all neurites normalized to mock control [one-way ANOVA, F (3, 302) = 9.658, p < 0.0001]. ***G,*** Bar graph quantification of β-actin intensity in primary neurites normalized to % mock control [one-way ANOVA, F (3, 171) = 4.695, p = 0.0035]. ***H,*** Bar graph quantification of β-actin intensity in secondary neurites normalized to mock control [one-way ANOVA, F (3, 127) = 3.227, p = 0.0248]. n = 3 independent experiments with n = 15-16 neurons per condition analyzed. ***I,*** Representative images immunostained for MAP2. ***J,*** Bar graph quantification of fluorescent MAP2 intensity in all neurites normalized to mock control [one-way ANOVA, F (3, 414) = 1.950, p = 0.1208]. ***K,*** Bar graph quantification of MAP2 intensity in primary neurites normalized to mock control [one-way ANOVA, F (3, 220) = 1.010, p = 0.3891]. ***L,*** Bar graph quantification of MAP2 intensity in secondary neurites normalized to mock control [one-way ANOVA, F (3, 191) = 1.652. p = 0.1788]. n = 3 independent experiments with n = 16-18 neurons per condition analyzed. Scale bars for ***A, E, I*** = 10 μm.

In contrast, we observed a 45% increase in β-actin intensity across all neurites after Reelin stimulation compared to mock control (*p* < 0.0001, Fig. 3*E, F*). Likewise, we observed a 37% increase in β-actin intensity in primary neurites *p* = 0.0059 and a 57% increase in secondary neurites, *p* = 0.0367 after Reelin stimulation compared to mock control (Fig. 3 *G, H*). However, β-actin intensity was not changed in the presence of GST-RAP or LY294002. Since we did not observe any changes in β-actin protein levels by immunoblotting after Reelin stimulation (Fig. 4*E, H*), it is likely that the increase in β-actin intensity seen by immunofluorescence after Reelin stimulation is an increase in monomeric and/or oligomeric β-actin due to F-actin remodeling and not a result of *de novo* translation of β-actin.

**Figure 4.**
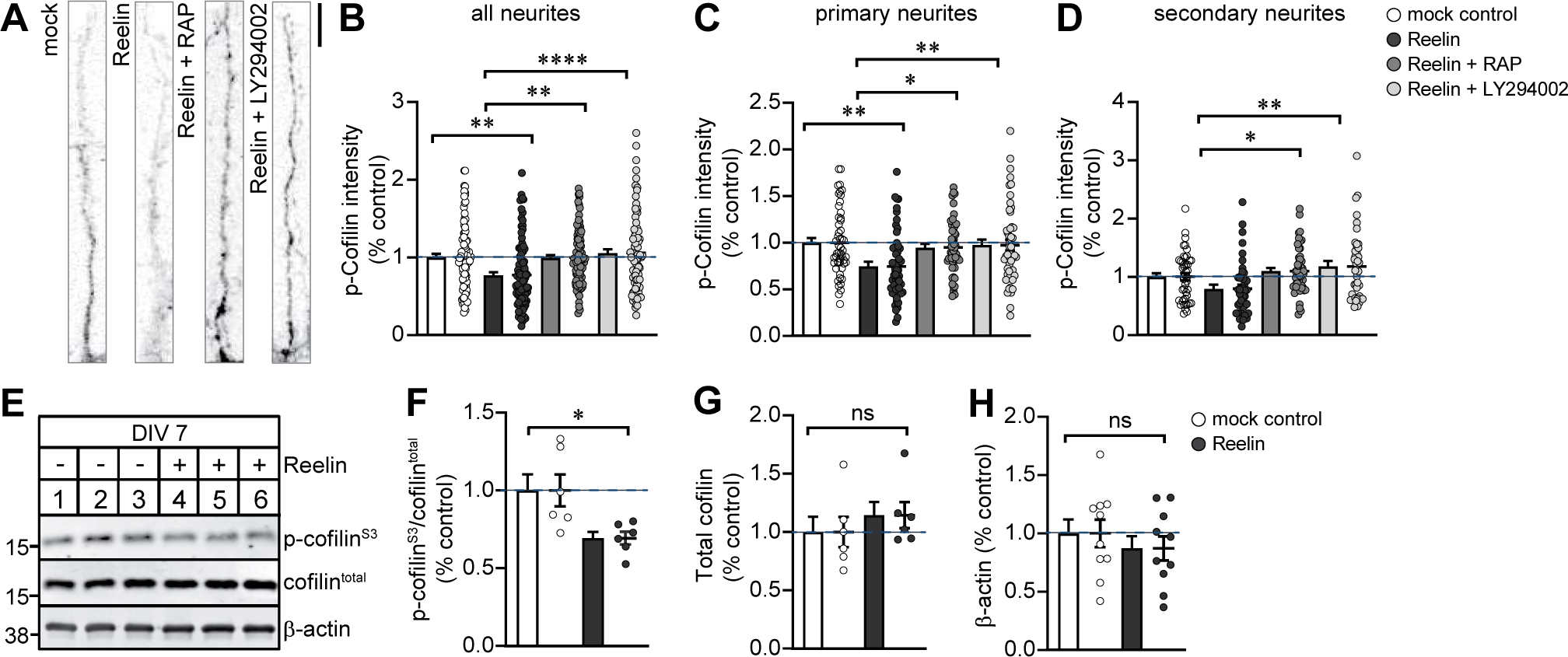
Reelin reduces n-cofilin phosphorylation at Ser3. ***A,*** Primary murine neurons at DIV 7 were treated with mock control, 50 nM Reelin, 100 nM GST-RAP for 2 h prior to Reelin or 50 μM LY294002 for 30 min prior to Reelin stimulation (30 min). Representative images immunostained for phospho-cofilin at Ser3 showed a decrease in phospho-cofilin in neurites following Reelin stimulation. Scale bar = 10 μm. ***B,*** Bar graph quantification of fluorescent phospho-cofilin intensity in all neurites normalized to mock control [one-way ANOVA, F (3, 383) = 8.602, p < 0.0001]. ***C,*** Bar graph quantification of phospho-cofilin intensity in primary neurites normalized to % mock control [one-way ANOVA, F (3, 203) = 5.540, p = 0.0011]. ***D,*** Bar graph quantification of phospho-cofilin intensity in secondary neurites normalized to mock control [one-way ANOVA, F (3, 176) = 5.500, p = 0.0012]. n = 3 independent experiments with n = 14-16 neurons per condition analyzed. ***E,*** Representative immunoblots from DIV 7 neuronal lysates treated with 50 nM Reelin or mock control for phospho-cofilin at Ser3, total n-cofilin and β-actin. ***F,*** Bar graph quantification for the ratio of phosphorylated cofilin at Ser3 with total cofilin from ***E*** shows a decrease in cofilin phosphorylation following Reelin stimulation [Unpaired *t*-test, t = 2.763, df = 10, *p* = 0.0200]. ***G-H,*** Bar graph quantification for total cofilin and β-actin, respectively. n = 6-10 individual replicates.

To determine whether the cytoskeleton changes we observed were specific to actin remodeling, we immunolabeled neurons against the microtubule associated protein, MAP2. We did not observe any MAP2 intensity changes across all four treatment conditions in all neurites observed (Fig. 3*I-L*), suggesting Reelin’s effect on the cytoskeleton was specific to actin remodeling at DIV 7. Together, the increase in β-actin intensity in conjunction with the decrease in phalloidin intensity demonstrates the canonical Reelin signaling modulates actin dynamics in nascent neurites, supporting the proteomic changes we observed related to regulation of cytoskeleton organization, particularly for actin filament-based processes.

### Reelin regulates n-cofilin phosphorylation at serine3 in nascent neurites

After observing that Reelin stimulation leads to a decrease in F-actin, we next asked whether the decrease is mediated through the actin severing protein, n-cofilin since it has been reported that Reelin signaling modulates serine3 phosphorylation of n-cofilin (Chai et al., 2009; Leemhuis et al., 2010). A decrease in phosphorylation at serine3 promotes n-cofilin to sever F-actin which allows for actin rearrangement in cell motility and navigation (Agnew et al., 1995; Carlier et al., 1997; Ghosh et al., 2004; Bravo-Cordero et al., 2013; Chen et al., 2015). To test this, we immunolabeled against phospho-cofilin at serine3 in primary murine neurons following Reelin stimulation for 30 min. We found that Reelin decreased phospho-serine3-cofilin intensity by 23% when compared to mock control (*p* = 0.0011, Fig. 4*A, B*). The effect of phospho-serine3-cofilin intensity after Reelin stimulation was abolished in neurons treated with GST-RAP and LY294002, indicating that Reelin’s effects on n-cofilin phosphorylation is dependent on the canonical Reelin pathway. In addition, we found a similar 25% decrease in phospho-serine3-cofilin intensity in primary neurites after Reelin stimulation compared to mock control (*p* = 0.0021, Fig. 4*C*). While not significant, we observed a 21% decrease in phospho-serine3-cofilin intensity between Reelin and mock in secondary neurites (*p* = 0.1692, Fig. 4*D*). We did; however, observed a significant 27.5% and 32.5 % decrease in phospho-serine3-cofilin intensity when we compared the Reelin treated group to neurons treated with GST-RAP (*p* = 0.0104) and LY294002 (*p* = 0.0012), respectively (Fig. 4*D*).

We also compared the levels of phosphorylated n-cofilin protein from neuronal lysates collected from DIV 7 primary neurons stimulated with Reelin by immunoblotting for both n-cofilin serine3 phosphorylation and total n-cofilin. Western blot analysis showed that Reelin significantly decreased the ratio of phosphorylated n-cofilin at serine3 to total cofilin by 31% (*p* = 0.02, lanes 4-6, Fig. 4*E, F*). However, we did not observe any effect of Reelin on total cofilin and β-actin protein levels (Fig. 4*G, H*). Taken together, our data suggests acute Reelin signaling through the canonical Reelin–PI3K pathway regulates n-cofilin phosphorylation at serine3 in nascent neurites in culture.

### Reelin stimulation leads to *de novo* translation of aldolase A and mobilizes aldolase A from the cytoskeleton

We identified aldolase A, a glycolytic enzyme and actin-binding protein, from the quantitative proteomic dataset as a novel effector of the Reelin signaling pathway that may contribute to actin remodeling changes. Aldolase A was within the regulation of anatomical structure and cell morphogenesis pathway identified by GSEA. Several supporting evidences suggests aldolase A may play a role in Reelin-mediated neurite growth. First, aldolase A is a well-known F-actin binding protein shown to regulate cell motility in non-neuronal cells (Tochio et al., 2010; Ritterson Lew and Tolan, 2013). Second, aldolase A is expressed in the brain during early developmental time points that are associated with cell migration, growth, and extensive cytoskeletal reorganization (Weber, 1965; Lebherz and Rutter, 1969; Penhoet et al., 1969). Third, previous studies have shown that aldolase A and n-cofilin compete for the same binding site on F-actin (Gizak et al., 2019), and the inhibition of the aldolase A and actin interaction leads to loss of F-actin stress fibers in a n-cofilin dependent manner (Gizak et al., 2019; Sun et al., 2021). Lastly, aldolase A is an effector of the insulin-PI3K pathway that regulates epithelial cell’s metabolism through mobilization of aldolase A from the actin cytoskeleton (Hu et al., 2016). Given that we showed Reelin regulates actin dynamics and n-cofilin phosphorylation through PI3K signaling pathway, we posit that aldolase A may serve as a molecular effector of Reelin-mediated neurite growth by regulating actin remodeling dynamics.

To confirm whether Reelin leads to changes in aldolase A protein level expression, neuronal lysates collected from DIV 7 primary neurons stimulated with 50 nM Reelin or mock for 30 min were immunoblotted for aldolase A and β-actin. Western blot analysis showed that Reelin caused a 22% increase in aldolase A levels when compared to mock control (*p* = 0.0342, Fig. 5*A, B*). The effect of aldolase A after Reelin stimulation was prevented in neurons treated with GST-RAP or LY294002 indicating the increase in aldolase A protein levels was mediated through the canonical Reelin-PI3K signaling pathway. To determine whether the increase in aldolase A protein levels was a result of *de novo* translation or reduced protein degradation, DIV 7 neurons were treated with 40 μM anisomycin, a translation inhibitor, for 45 min prior to Reelin stimulation. Western blot analysis showed that Reelin caused a 46.5% increase in aldolase A levels when compared to mock control (*p* = 0.0499, Fig. 5*C, D*); however, this was abolished with anisomycin treatment demonstrating that the increase in aldolase A expression after Reelin stimulation is due to *de novo* translation. It is important to note that quantitative proteomics is typically based on the measurement of multiple spectra and peptides per protein that is accompanied by statistical confidence measures for each peptide, thus it has higher sensitivity of detecting protein changes compared to Western blotting approaches.

**Figure 5.**
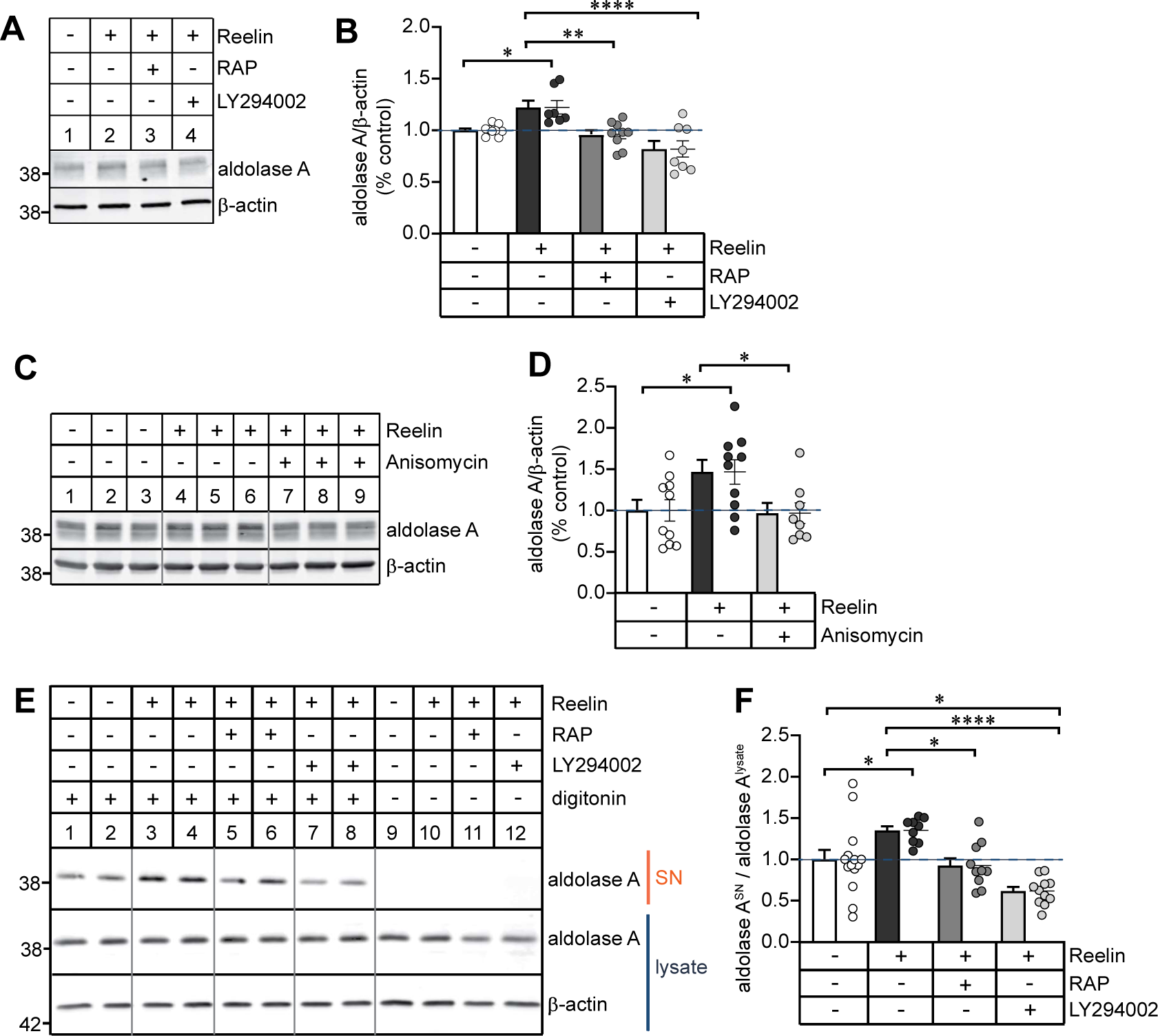
Aldolase A undergoes *de novo* translation and mobilizes from the cytoskeleton in response to Reelin stimulation. ***A,*** Representative immunoblots of DIV 7 neuronal lysates treated with mock control, 50 nM Reelin, 100 nM GST-RAP for 2 h prior to Reelin or 50 μM LY294002 for 30 min prior to Reelin stimulation for aldolase A and β-actin. ***B,*** Bar graph quantification of aldolase A normalized to mock control showed an increase in aldolase A protein levels after Reelin stimulation [one-way ANOVA, F (3, 29) = 8.998, p = 0.0002]. n = 7-9 individual replicates. ***C,*** Representative immunoblots of DIV 7 neuronal lysates treated with mock control (lanes 1-3), 50 nM Reelin (lanes 4-6) or 40 μM anisomycin for 45 min prior to Reelin stimulation (lanes 7-9) for aldolase A and β-actin. ***D,*** Bar graph quantification of aldolase A normalized to mock control showed an increase in aldolase A protein levels after Reelin stimulation but not in anisomycin-treated neurons [one-way ANOVA, F (2, 25) = 4.275, p = 0.0253]. n = 8-10 individual replicates. ***E,*** Representative immunoblots of DIV 7 neuronal lysates treated with mock control, 50 nM Reelin, 100 nM GST-RAP for 2 h prior to Reelin or 50 μM LY294002 for 30 min prior to Reelin stimulation and subject to digitonin permeabilization (lanes 1-8). Supernatant (SN) and cell lysate were subject to immunoblotting for aldolase A and β-actin as indicated. ***F,*** Quantification of aldolase A in the diffusible fraction (supernatant) to immobile (cell lysate) fraction show an increase in aldolase A in response to Reelin stimulation [one-way ANOVA, F (3, 41) = 10.45, p < 0.0001]. n = 9-15 individual replicates.

Previous studies had shown that aldolase A is mobilized from the actin cytoskeleton through PI3K activation in epithelial cells (Hu et al., 2016). To determine whether aldolase A is mobilized from the actin cytoskeleton through Reelin-PI3K signaling in primary murine neurons, we performed a digitonin permeabilization assay to allow for efflux of diffusible aldolase A and estimate the fraction of aldolase A in the soluble versus immobilized state based on separate collection of supernatant and cell lysates, respectively. DIV 7 neurons were treated with either mock, Reelin, RAP or LY294002 prior to Reelin stimulation and subsequently permeabilized with digitonin for 5 min at 4°C. We found a 35% increase in aldolase A in the supernatant fraction induced by Reelin stimulation indicating an increase in mobilized aldolase A when compared to mock control (*p* = 0.0337, lanes 3-4, Fig. 5*E, F*). Both RAP and LY294002 prevented the Reelin mediated effect on aldolase A mobilization. In fact, LY294002 reduced aldolase A mobilization by 39% compared to mock control (*p* = 0.0111, lanes 7-8, Fig. 5*E, F*). We did not detect any aldolase A in the supernatant from neurons that were not permeabilized with digitonin which served as a negative control for the assay (lanes 9-12, Fig. 5*E*). In summary, these results indicate that the effects of Reelin on aldolase A levels and mobilization from the cytoskeleton are dependent on the activation of PI3K.

### Aldolase A functions downstream of Reelin signaling in regulating dendrite growth

To directly test whether aldolase A functions downstream of Reelin signaling, we infected primary murine neurons with lentiviral *Aldoa* shRNA at DIV 1 to determine the effects of aldolase A knockdown on dendritic development following Reelin treatment. We found that *Aldoa* shRNA efficiently downregulated aldolase A protein expression at DIV 7 and resulted in a significant 64% knockdown compared to *scrambled* (*scr*) control shRNA (*p* = 0.0179, lane 2, Fig. 6*A, B*). In contrast, infection of lentiviral *Aldoa* shRNA in neurons had no effect on β-actin levels (Fig. 6*A, B*). Since aldolase A is one of two aldolase isozymes that is expressed in the brain, we also immunoblotted for aldolase C and found no difference in aldolase C levels demonstrating that *Aldoa* shRNA is specific to aldolase A (Fig. 6*A, B*).

**Figure 6.**
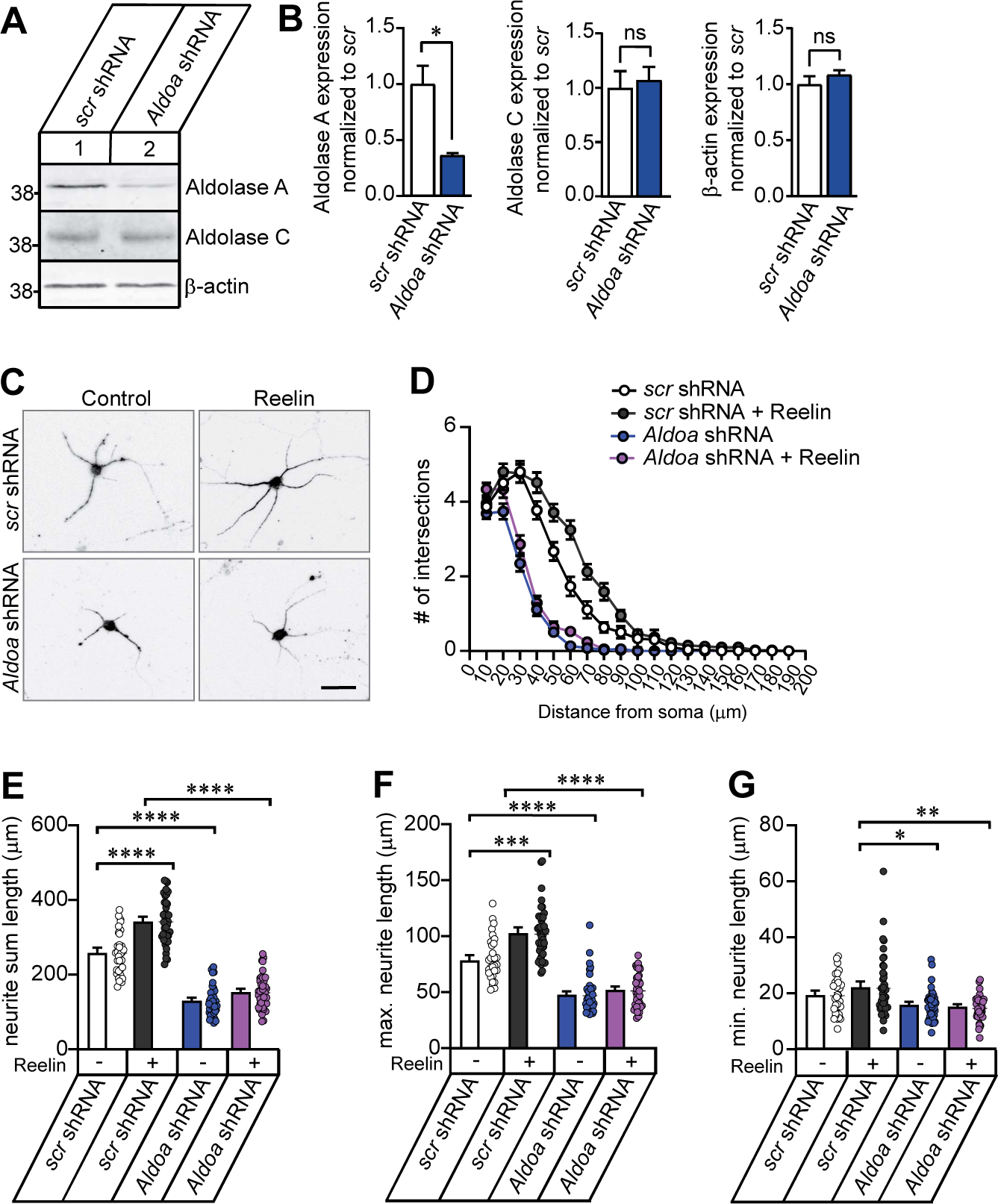
Aldolase A is necessary for Reelin-mediated neurite outgrowth and arborization. Primary murine neurons on DIV 1 were infected with *scr* or *Aldoa* shRNA and analyzed at DIV 7. ***A,*** Representative immunoblots showing aldolase A and aldolase C levels from DIV 7 neuronal lysate after infection with *scr* or *Aldoa* shRNA. β-actin served as loading control. ***B,*** Bar graph quantifications of aldolase A, aldolase C, and β-actin expression normalized to *scr* shRNA showed decreased protein level specifically for aldolase A [Unpaired *t-test*, t = 3.874, df = 4, *p* = 0.0179]. n = 3 individual replicates. ***C,*** Representative images of DIV 7 neurons infected with *scr* or *Aldoa* shRNA and stimulated with either mock control or 50 nM Reelin once a day for 72 h starting at DIV 4 and immunostained against MAP2 at DIV 7. Scale bar = 12.5 μm. ***D,*** Sholl analysis comparing neuronal arborization complexity between *scr* or *Aldoa* shRNA with mock and Reelin treatment showed reduced dendritic growth and arborization in aldolase A knockdown neurons and Reelin stimulation had no effect in neurons following aldolase A knockdown [two-way ANOVA, F (54, 2793) = 21.80, *p* < 0.0001]. n = 3 independent experiments with n = 30-42 cells per condition analyzed. ***E,*** Bar graph quantification of the average total neurite sum length [one-way ANOVA, F (3, 157) = 89.41, *p* < 0.0001]. ***F,*** Bar graph quantification of the average maximum neurite sum length [one-way ANOVA, F (3, 158) = 46.06, *p* < 0.0001]. ***G,*** Bar graph quantification of the average minimum neurite length [one-way ANOVA, F (3, 158) = 5.149, *p* = 0.0020]. For ***E-G***, n = 3 independent experiments with n = 32-44 cells per condition analyzed.

To investigate whether aldolase A functions downstream of Reelin signaling on dendritic branching, primary murine neurons were infected at DIV 1 with *scr* or *Aldoa* shRNA and starting at DIV 4, neurons were treated with 50 nM Reelin or mock control once a day for 72 h and immunolabeled against dendritic marker, MAP2 at DIV 7. We performed Sholl analysis to examine global changes in neuronal morphology, specifically the complexity of the dendritic arbor and the overall pattern of arborization. Sholl analysis revealed that shRNA knockdown of aldolase A greatly reduced dendritic growth where it shifted the distribution downward and to the left compared with *scr* shRNA neurons, indicating both a decrease in the number and length of the neurites (Fig. 6*C, D*). The average total neurite length was 49.5% shorter in neurons infected with *Aldoa* shRNA compared to *scr* shRNA infected neurons (*p* < 0.0001, Fig. 6*E*). Similarly, the average maximum neurite length was decreased by 40% in *Aldoa* shRNA infected neurons (*p* < 0.0001, Fig. 6*F*), but had no effect on the average minimum neurite length between *scr* and *Aldoa* shRNA infected neurons (Fig. 6*G*). Meanwhile, Reelin induced a 32% increase in the average total neurite length in *scr* shRNA neurons compared to mock control (*p* < 0.0001, Fig. 6*E*). Similarly, the average maximum length of the longest neurite was increased by 31% following Reelin stimulation (*p* = 0.0002, Fig. 6*F*), but had no effect in the average minimum neurite length (Fig. 6*G*). Interestingly, Reelin stimulation had no effect in neurons following aldolase A knockdown, indicating that aldolase A lies downstream of Reelin signaling and is necessary for the effects of Reelin on dendritic outgrowth (Fig. 6*C-G*).

### Aldolase A knockdown disrupts neuronal migration and morphology *in vivo*

Since we found aldolase A functions downstream of the Reelin signaling pathway, we next asked whether aldolase A knockdown impacts neuronal migration and neuronal architecture *in vivo*. To test this, we performed *in utero* electroporation with GFP-tagged *scr* or *Aldoa* shRNA on mouse embryos at E15.5 and analyzed both the position of GFP positive cells within the cortical layers and morphology located in primary somatosensory cortex at P14 to account for uniform distribution of cells along the rostro-caudal axis. As expected, we found most GFP-positive neurons electroporated with *scr* shRNA were in upper layers II and III that expressed Satb2 which we used as a marker to distinguish cortical layers for analysis (Fig. 7*A, B*). Interestingly, we found neurons electroporated with *Aldoa* shRNA distributed deeper within layers II and III. We, therefore, measured the distance of the soma from the pia and compared neurons electroporated with *scr* and *Aldoa* shRNA. We found neurons electroporated with *Aldoa* shRNA were 57.75 μm deeper in the cortex compared to *scr* electroporated neurons (p < 0.0001, Fig. 7*C*) suggesting that aldolase A may play a role in neuronal migration.

**Figure 7.**
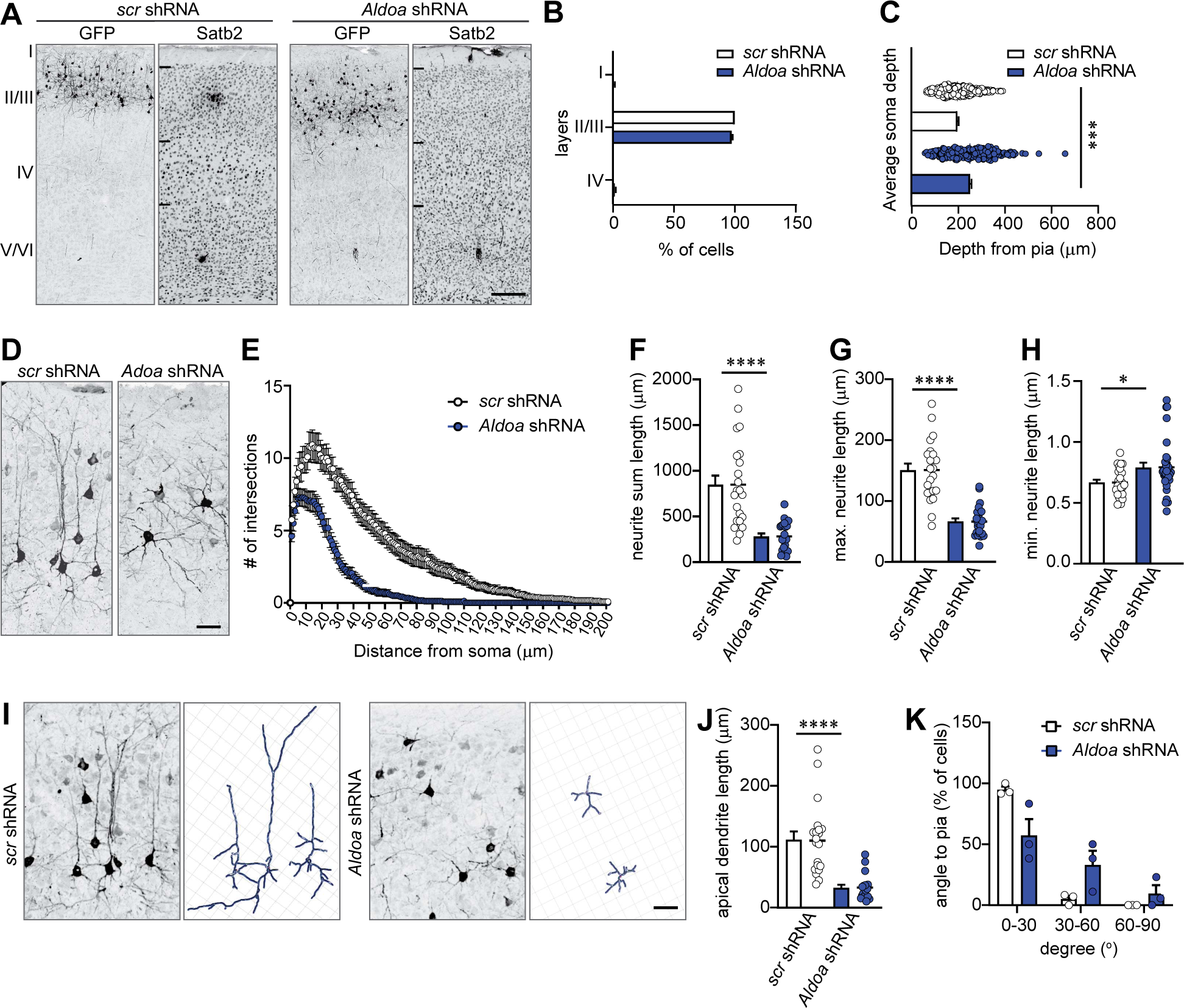
Aldolase A is required for neuronal arborization and apical dendrite polarity in the mouse cortex. Mouse embryos were electroporated *in utero* with GFP-tagged *scr* control or *Aldoa* shRNA at E15.5 and analyzed at P14. ***A,*** Representative images of brain sections electroporated with GFP-tagged *scr* control or *Aldoa* shRNA and immunostained against GFP and Satb2 to visualize electroporated cells and marker to distinguish cortical layers in somatosensory S1 cortex, respectively. Scale bar = 175 μm. ***B,*** Bar graph quantification showing the percentage of GFP-positive neurons across cortical layers between *scr* and *Aldoa* shRNA electroporated neurons. n = 3 mice per group with n = 402-639 cells per group analyzed. ***C,*** Bar graph quantification showing the average soma depth of GFP-positive electroporated neurons from the pia showed aldolase A knockdown neurons reside deeper in the cortex compared to *scr* electroporated neurons [Unpaired *t*-test, t = 7.889, df = 556, *p* < 0.0001]. n = 3 brains per group with n = 264-294 cells per group analyzed. ***D,*** Representative images of GFP-positive neurons in layers II/III of S1 from brains electroporated at E15.5 with *scr* or *Aldoa* shRNA showing the disrupted morphology and arborization of aldolase A knockdown neurons compared to *scr* control. Scale bar = 25 μm. ***E,*** Sholl Analysis showed aldolase A knockdown neurons have shorter total neurite length and less neuronal arborization compared to *scr* control neurons. A two-way ANOVA was used to determine the statistical significance of the interaction between the length of the neurites and number of crossings at each radius for each treatment condition [two-way ANOVA F (222, 14718) = 14.06, *p* < 0.0001]. n = 3 brains per condition and n = 31-37 neurons per condition analyzed. ***F,*** Bar graph quantification showing aldolase A knockdown neurons led to a decrease in the average neurite sum length compared to *scr* control electroporated neurons [Unpaired *t*-test, t = 6.478, df = 63, *p* < 0.0001]. ***G,*** Bar graph quantification of the average maximum neurite length between *scr* and *Aldoa* shRNA electroporated neurons [Unpaired *t*-test, t = 7.272, df = 62, *p* < 0.0001]. ***H,*** Bar graph quantification of the average minimum neurite length between *scr* and *Aldoa* shRNA electroporated neurons [Unpaired *t*-test, t = 2.559, df = 55, *p* = 0.0133]. For ***F-H***, n = 3 brains per group with n = 25-35 neurons analyzed. ***I,*** Representative images of GFP-positive neurons in layers II/III of S1 from brains electroporated with either *scr* or *Aldoa* shRNA (left panel) and IMARIS generated 3D wire structure reconstruction of the same neurons (right panel) showing their morphology and apical neurite orientation and length is altered in aldolase A knockdown neurons. Scale bar = 25 μm. ***J,*** Bar graph quantification showing a reduction in the average length of apical neurites for aldolase A knockdown neurons [Unpaired *t*-test, t = 6.164, df = 65, *p* < 0.0001]. n = 3 brains per group and n = 33-34 neurons analyzed. ***K,*** Percentage of GFP-positive neurons with an apical dendrite oriented between 0-30°, 31-60° and 61-90° to the pia were quantified. Quantification revealed control electroporated neurons have apical dendrites oriented mostly within 0-30° compared to aldolase A knockdown neurons [two-way ANOVA, F (2, 12) = 9.129, *p* = 0.0039]. n = 3 brains per group with n = 30-35 neurons analyzed.

We next examined whether aldolase A is necessary for proper neuronal growth and arborization and performed Sholl analysis on neurons electroporated with *scr* or *Aldoa* shRNA. Sholl analysis revealed aldolase A knockdown neurons exhibited a significant reduction in both dendritic length and arbor complexity compared to *scr* electroporated neurons (*p* < 0.0001, Fig. 7*D, E*). The average total neurite length was 65.6% shorter in aldolase A knockdown neurons compared to *scr* electroporated neurons (*p* < 0.0001, Fig. 7*F*). Similarly, the average maximum neurite length was decreased by 53.4% in *Aldoa* shRNA electroporated neurons (*p* < 0.0001, Fig. 7*G*). Interestingly, we found a small and significant increase in the average minimum neurite length between *scr* and *Aldoa* shRNA electroporated neurons (*p* = 0.0133, Fig. 7*H*).

Not only did aldolase A knockdown neurons have a significant decrease in neurite length and arborization, it was apparent they also lacked the apical-basal polarity typical of cortical pyramidal neurons (Fig. 7*I*). As such, we measured the length and orientation of the apical dendrite in GFP-positive neurons of *scr* control and *Aldoa* shRNA electroporated neurons at P14. Our results demonstrate that aldolase A knockdown caused a 69.24% decrease in the length of the apical dendrite when compared to controls (*p* < 0.0001, Fig. 7*J*). We examined apical dendrite orientation and calculated this as an angle in relation to the pial surface and binned the data into three groups: 0-30°, 31-60° and 61-90° from the pial surface. Nearly 94.83% of GFP-positive neurons electroporated with *scr* control had their apical dendrite oriented between 0-30° to the pia. However, only 57.26% of aldolase A knockdown neuron kept this normal, apical orientation (*p* = 0.017, Fig. 7*K*). Instead, we found nearly 33.2% of aldolase A knockdown neurons had an apical dendrite oriented between 31-60° and 9.55% had their apical dendrite between 61-90°, both trending higher when compared to *scr* control neurons. Altogether, these data support the conclusion that aldolase A is necessary for proper neuron arborization and is, particularly important for the growth and orientation of the apical dendrite of pyramidal neurons in the mouse cortex.

### Aldolase A requires actin binding activity for proper neuronal arborization

Because aldolase A serves as both a glycolytic enzyme and actin binding protein, it is in a unique position to regulate the metabolic demands and the structural rearrangements of the cytoskeleton necessary for the energy intensive process of neurite growth and arborization. To determine which functional activity of aldolase A is required for proper neuronal arborization, we examined two aldolase variants defective in either glycolytic (D33S) or actin-binding activity (R42A) that have been previously characterized in non-neuronal cells (Wang et al., 1996; Ritterson Lew and Tolan, 2012, 2013; Hu et al., 2016). The D33S aldolase A variant is catalytically dead but retains F-actin binding activity; whereas the R42A variant abolishes the F-actin binding activity but retains glycolytic activity. We generated three lentiviral aldolase A constructs: wild type, D33S and R42A and we titered each lentivirus uniformly to perform rescue experiments of aldolase A’s knockdown effects on dendritic arborization. Primary murine neurons were infected on DIV 1 and we performed immunolabeling against MAP2 at DIV 7. We compared the effects of *scr* shRNA, *Aldoa* shRNA, *Aldoa* shRNA rescued with mouse aldolase A wild type, catalytically dead D33S or actin-binding deficient R42A variant. The *Aldoa* shRNA-mediated knockdown of aldolase A decreased neurite arborization compared to *scr* shRNA (*p* < 0.0001, Fig. 8*A, B*). Both aldolase A wild type and catalytically dead D33S rescued the *Aldoa* shRNA phenotype, whereas the actin-binding deficient R42A was unable to rescue the neurite arborization effects of aldolase A knockdown on dendritic outgrowth (Fig. 8*A, B*).

**Figure 8.**
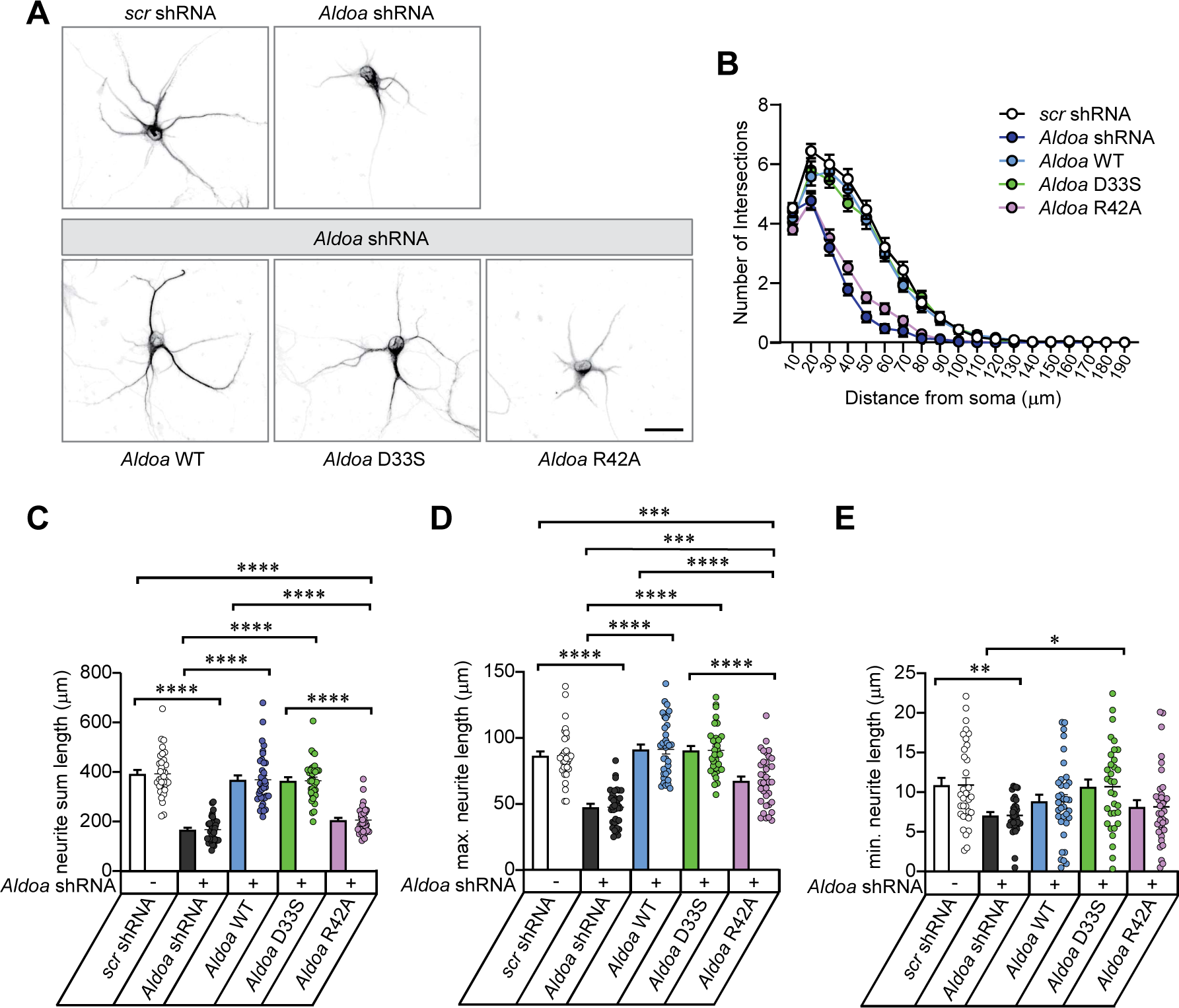
Aldolase A requires actin binding function to regulate proper neuronal arborization. ***A,*** Representative images of DIV 7 neurons immunostained against MAP2 infected with *scr* shRNA, *Aldoa* shRNA, or *Aldoa* shRNA rescue with aldolase A wild type (WT), D33S, or R42A variant on DIV1. Scale bar = 25 µm. ***B,*** Sholl analysis showed *Aldoa* shRNA infected neurons exhibit reduced dendritic length and arborization compared to *scr* control. Lentiviral infection with aldolase WT or D33S variant rescued *Aldoa* shRNA induced phenotype while R42A variant was unable to rescue the phenotype. Two-Way ANOVA analysis used to determine statistical significance of the interaction between neurite length and number of intersections at each radius for each condition [two-way ANOVA, F (72, 3230) = 14.70, *p* < 0.0001]. n = 3 independent experiments with n = 34-36 cells analyzed per condition. ***C,*** Bar graph quantification of the average total neurite sum length One-Way ANOVA used to determine statistical significance [one-way ANOVA F (4, 170) = 63.91, *p* < 0.0001]. ***D,*** Bar graph quantification of the average maximum neurite sum length [one-way ANOVA F (4, 168) = 36.06, *p* < 0.0001]. ***E,*** Bar graph quantification of the average minimum neurite length [one-way ANOVA F (4, 165) = 4.176, *p* = 0.003]. For ***C-E***, n = 3 independent experiments with n = 33-36 neurons analyzed per condition.

We systematically measured the average total, maximum and minimum neurite length between the conditions. The average total neurite length was 57.4% shorter in neurons infected with *Aldoa* shRNA compared to *scr*-infected neurons (*p* < 0.0001, Fig. 8*C*). Lentiviral infection with aldolase A wild type and catalytically dead D33S variant fully rescued the effects of aldolase A knockdown phenotype (*p* < 0.0001, Fig. 8*C*). In contrast, the actin-binding deficient R42A variant led to a 47.8% reduction in average total neurite length compared to *scr* shRNA and was unable to rescue the aldolase A knockdown phenotype (*p* < 0.0001, Fig. 8*C*). Similarly, the average maximum neurite length was decreased by 44.9% in *Aldoa* shRNA infected neurons (*p* < 0.0001, Fig. 8*D*). Both aldolase A wild type and D33S fully rescued the *Aldoa* shRNA phenotype (*p* < 0.0001, Fig. 8*D*); however, the R42A variant was unable to rescue the maximum neurite length following aldolase A knockdown when compared to *scr* shRNA control (*p* = 0.0002, Fig. 8*D*). Likewise, the average minimum length of the shortest neurite was rescued by the catalytically dead D33S aldolase A variant (*p* = 0.0152, Fig. 8*E*). Together, these results provide strong evidence that the actin binding function of aldolase A is a key regulator controlling dendritic outgrowth.

## Discussion

In this study, we showed Reelin selectively activates S6K1 phosphorylation through the PI3K/Akt and mTORC1 complex pathways during initial stages axonal and dendritic outgrowth, a process that differs in mature neurons in culture (Fig. 1). This aligns with previous studies that demonstrated Reelin signaling to PI3K/Akt is necessary for Reelin-induced neurite outgrowth (Beffert et al., 2002; Jossin and Goffinet, 2007; Leemhuis et al., 2010). Through an unbiased proteomics analysis to delineate between canonical and non-canonical Reelin signaling during dendritic outgrowth in primary murine neurons, we found significant crosstalk related to cytoskeleton regulation, neuron projection development and actin filament-based processes (Fig. 2). Notably, gene sets influenced by the non-canonical Reelin pathway include protein translation, mRNA metabolic processes and ribonucleoprotein complex biogenesis, highlighting distinct impacts on neuronal structure and function.

We explored how Reelin specifically influences actin dynamics in dendritic development (Fig. 3). We showed that Reelin stimulation reduces F-actin intensity while increasing total β-actin intensity in developing neurites. However, this increase in β-actin was not corroborated by Western blot analysis, suggesting the changes we observed in immunocytochemistry might be due to an increase in monomeric actin following F-actin depolymerization which will need to further examined. Accompanying these changes was a reduction in cofilin serine3 phosphorylation (Fig. 4), indicating enhanced cofilin severing activity. This suggests that Reelin promotes cofilin-mediated actin depolymerization and rearrangement in neurites *in vitro*, in contrast to previous studies that indicated Reelin signaling through PI3K increased cofilin serine3 phosphorylation, stabilizing the actin cytoskeleton in the leading processes of migrating neurons during embryonic mouse brain development (Chai et al., 2009; Frotscher et al., 2017). We posit Reelin’s dual role in modulating cofilin activity, either inhibiting or promoting cofilin severing activity, depending on the developmental stage. Reelin inhibits cofilin severing activity and stabilize the actin cytoskeleton to act as a stop signal for migrating neurons which is required for orientation and directed migration of cortical neurons (Chai et al., 2009), but it could also function to promote cofilin severing activity by decreasing cofilin serine3 phosphorylation to allow for actin rearrangements essential for neurite outgrowth and arborization.

Reelin is a commanding regulator on dendritic growth during different stages of brain development, particularly in embryonic and postnatal stages. Previous studies have shown that dendritic growth of pyramidal cells is disrupted in both homozygous and heterozygous *reeler* mice (Pinto Lord and Caviness, 1979; Niu et al., 2004). Addition of Reelin *in vitro* has been shown to increase dendritic growth of hippocampal neurons in both wild type and *reeler* mutant mice (Niu et al., 2004; Jossin and Goffinet, 2007; Matsuki et al., 2008). This enhancement of dendritic growth by Reelin is evident during early brain development (Nichols and Olson, 2010; Kupferman et al., 2014; Chai et al., 2015; Kohno et al., 2015; O’Dell et al., 2015). Our current study aligns with these findings, focusing on the *in vitro* effects of Reelin on the dendritic growth and branching of embryonic hippocampal neurons. However, Reelin exerts an opposing effect on dendritic growth in cortical pyramidal neurons and forebrain interneurons during postnatal development (Yabut et al., 2007; Chameau et al., 2009; Hamad et al., 2021). Specifically, the N-terminal fragment of Reelin has been shown to control and restrict the postnatal maturation of apical dendrites in pyramidal cortical neurons, a process mediated by integrin receptors (Chameau et al., 2009). This suggests that the postnatal effects of Reelin might be attributed to the differential impact of its proteolytic fragments interacting with non-canonical receptors. Our work contributes to the growing body of evidence on the role of Reelin in dendritic development. We provide proteomic changes into how both canonical and non-canonical Reelin pathways regulate dendrite growth in embryonic cultured neurons. However, it remains unclear whether these *in vitro* observations fully reflect the *in vivo* mechanisms and effects of Reelin on dendritic development across various stages of brain maturation.

Our proteomics screen identified aldolase A, a glycolytic enzyme and actin-binding protein, as a critical component in Reelin-mediated actin dynamics (Arnold and Pette, 1968; Arnold et al., 1971; Wang et al., 1996; Kusakabe et al., 1997; Jewett and Sibley, 2003). We demonstrated that Reelin stimulates *de novo* translation of aldolase A, mobilizing soluble aldolase A from the cytoskeleton, linking aldolase A directly to Reelin signaling and its novel role in neuronal development (Fig. 5). We show downregulation of aldolase A in primary murine neurons led to shorter, less branched dendrites, and Reelin treatment was ineffective following aldolase A knockdown, indicating aldolase A is necessary for Reelin’s effects on dendritic growth. Similar results were observed in cortical neurons in the developing mouse brain, with impaired neuronal migration, aberrant length, arborization and orientation of the apical dendrite (Fig. 7), reinforcing the essential role of aldolase A in proper neuron development.

We further investigated aldolase A’s dual functions in neurite outgrowth and demonstrated that the actin-binding ability of aldolase A, rather than its glycolytic function, is critical for neurite development using site-directed variants of aldolase A (Fig. 8). As previously characterized, the D33S aldolase variant is catalytically dead and retains F-actin binding ability while the R42A mutant is catalytically active and unable to bind F-actin (Wang et al., 1996; Ritterson Lew and Tolan, 2013; Hu et al., 2016). We demonstrated that both the wild type and catalytically inactive D33S aldolase variant, which retains F-actin binding ability, rescued the effects of *Aldoa* shRNA knockdown phenotype in primary neurons, while the R42A mutant, unable to bind F-actin, showed impaired neurite growth and arborization. This finding reinforces aldolase A’s role in cell growth and motility, independent of its glycolytic activity, in non-neuronal cells as observed in previous studies (Tochio et al., 2010; Ritterson Lew and Tolan, 2012, 2013; Gizak et al., 2019). Our data provides a direct link between aldolase A and neuronal development downstream of Reelin signaling, establishing a new functional role for aldolase A in neurons and expanding our understanding of how Reelin regulates actin filament dynamics related to dendrite growth.

In summary, our data elucidates how Reelin canonical and non-canonical signaling pathways uniquely affect neuronal structure and function *in vitro*. We demonstrate Reelin’s acute regulation of the actin cytoskeleton, highlighting aldolase A’s pivotal role as an actin binding protein which establishes an unrecognized function of aldolase A in the brain. We propose a model where Reelin modulates cofilin serine3 phosphorylation, facilitating aldolase A’s mobilization and enabling cofilin-dependent actin depolymerization and rearrangement essential for neurite extension and arborization.

## Conflict of interest

The authors declare no competing financial interests.

## Acknowledgments

We would like to thank Dr. Dean Tolan who provided suggestions for this work. This work was supported by the National Institute of Aging Grants AG059762 and the Harold and Margaret Southerland Alzheimer’s Research Fund to U.B. and A.H.

## Author contributions

G.D.L., U.B., A.H. designed research; G.D.L., S.N., N.L. performed research; G.D.L., W.L., S.N., N.L. analyzed data; G.D.L., A.E., U.B., and A.H. provided supervision; G.D.L., U.B., and A.H. wrote the paper.

## Figure Legends

**Supplementary Figure 1.**
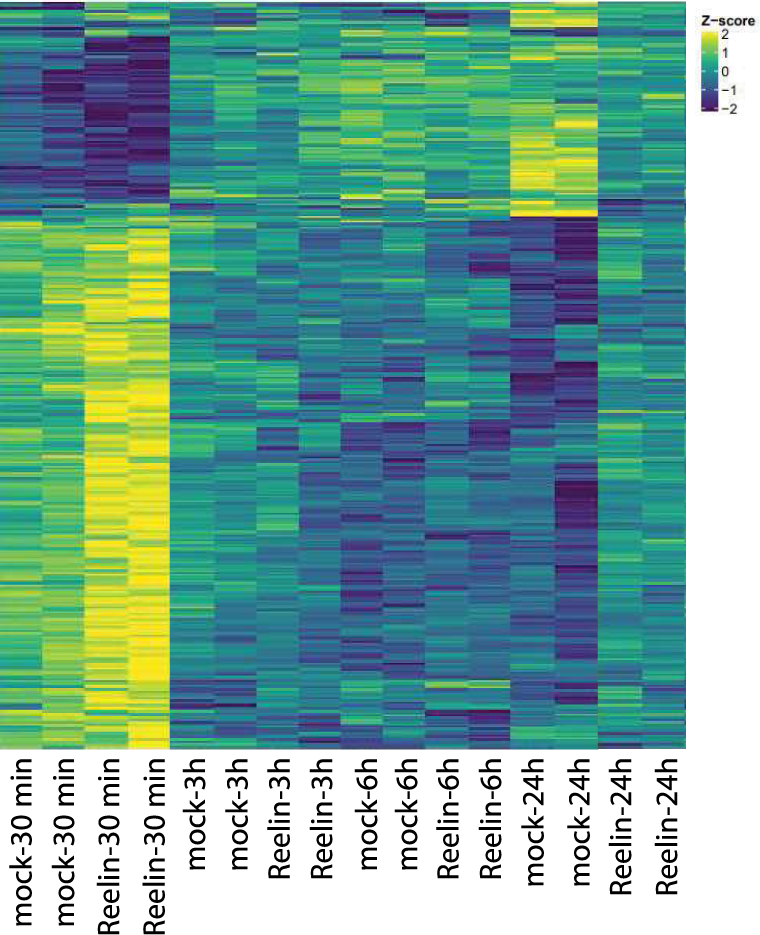
Label free quantification proteomics analysis of Reelin time-course treatment on wild type neurons at DIV 7. Each column represents an individual sample with duplicate treatments grouped together. Heatmap showing differentially expressed features identified with an FDR < 0.05. Feature intensity is represented as a Z-score between +2 (Yellow) and -2 (Blue), higher Z-scores correspond to greater relative expression while lower Z-scores correspond to lower relative expression. Dendrogram clustering groups treatment conditions by the similarity of the proteomic profile

**Supplementary Figure 2.**
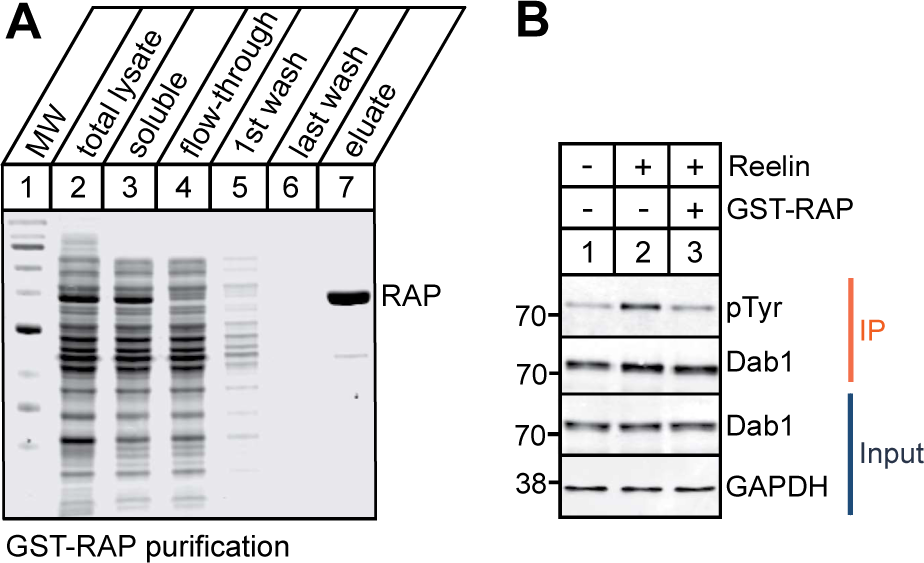
GST-RAP inhibits Dab1 phosphorylation. ***A,*** Coomassie showing fractions taken from GST-RAP production and purification. ***B,*** Representative immunoblot of tyrosine-phosphorylated Dab1 immunoprecipitated from DIV 7 primary murine neurons demonstrating GST-RAP prevented Reelin-induced Dab1 phosphorylation. Neurons were treated with 50 nM Reelin or mock control for 30 min or 100 nM GST-RAP for 2 h prior to 30 min Reelin treatment.s

## Notes

### Competing Interest Statement

The authors have declared no competing interest.

